# The formation and propagation of human Robertsonian chromosomes

**DOI:** 10.1101/2024.09.24.614821

**Authors:** Leonardo Gomes de Lima, Andrea Guarracino, Sergey Koren, Tamara Potapova, Sean McKinney, Arang Rhie, Steven J Solar, Chris Seidel, Brandon Fagen, Brian P Walenz, Gerard G Bouffard, Shelise Y Brooks, Michael Peterson, Kate Hall, Juyun Crawford, Alice C Young, Brandon D Pickett, Erik Garrison, Adam M Phillippy, Jennifer L. Gerton

## Abstract

Robertsonian chromosomes are a type of variant chromosome found commonly in nature. Present in one in 800 humans, these chromosomes can underlie infertility, trisomies, and increased cancer incidence. Recognized cytogenetically for more than a century, their origins have remained mysterious. Recent advances in genomics allowed us to assemble three human Robertsonian chromosomes completely. We identify a common breakpoint and epigenetic changes in centromeres that provide insight into the formation and propagation of common Robertsonian translocations. Further investigation of the assembled genomes of chimpanzee and bonobo highlights the structural features of the human genome that uniquely enable the specific crossover event that creates these chromosomes. Resolving the structure and epigenetic features of human Robertsonian chromosomes at a molecular level paves the way to understanding how chromosomal structural variation occurs more generally, and how chromosomes evolve.

## Introduction

Robertsonian chromosomes (ROBs) are structurally variant chromosomes created by the fusion of two telo- or acrocentric chromosomes to make a single metacentric chromosome. First recognized in 1916 by Robertson in grasshopper karyotypes ^1^, these fusions are a common occurrence in nature, having since been recognized in many branches of life including plants ^2^, vertebrates ^3^ ^4^, and invertebrates ^5^. Robertsonian fusions (or translocations) are the most common karyotypic change in mammals ^6^. ROBs create challenges for meiosis, potentially leading to subfertility, reproductive isolation, and speciation ^7,8^. In humans, ROBs contribute to trisomies like Down and Patau syndrome ^9^, and are associated with elevated rates of certain cancers ^10^ and uniparental disomy ^11^. Despite their frequent occurrence and significant impacts on fertility, speciation, and human health, the underlying mechanisms and evolutionary origins that explain why these chromosomes form so often in nature remain unknown.

In humans, ROBs occur in 1 out of 800 births ^12–14^, most commonly in female meiosis ^15,16^. The most common ROBs are the fusion of acrocentric chromosome 14 with either chromosome 13 or chromosome 21, suspected to form by a similar specific mechanism ^17^. In these fusions, the long arms are joined and parts of the short arms are lost. ROBs can occur de novo or be inherited ^18^. The recent complete assembly of the human CHM13 genome ^19^, including the short arms of the acrocentric chromosomes, followed by a population-level analysis of these regions for the first time ^20^, revealed Mb-sized homology blocks shared between acrocentric chromosomes, termed pseudo-homologous regions (PHRs). The existence of PHRs implies frequent interhomolog recombination and a working model for how ROBs form ^21^.

Herein we fully assemble three common ROBs: two 13;14 fusions and one 14;21 fusion. In all three cases, the ROBs have two centromere arrays, have lost all ribosomal 45S DNA repeats (or nucleolar organizer regions (NORs)), and are fused near a macrosatellite array composed of SST1 repeats ^22,23^. We demonstrate that the SST1 arrays on NOR-containing chromosomes, previously also called NBL2 ^24^, bear hallmarks of exchange between chromosomes, including enrichment for PRDM9 DNA binding motifs which are associated with recombination hotspot activity ^25^. Further analysis of SST1 repeats and segmental duplications in human and nonhuman primate genomes suggests SST1 may have a broader role in genome rearrangements and evolution beyond Robertsonian fusions. Additionally, analysis of the two centromeric arrays on each ROB indicates epigenetic adaptations to support stable mitotic transmission, consistent with previous observations^26,27^. In conclusion, we provide the first complete assemblies of common ROBs, precise mapping of their fusion sites, and novel insight into their formation and transmission mechanisms.

## Results

A working model for the formation of common ROBs postulated that they occur due to 1) sequence homology between nonhomologous chromosomes, provided by the PHRs, 2) proximity of the PHRs, provided by co-location of rDNA arrays from different chromosomes in the nucleolus, 3) recombination initiation in meiosis to form a crossover, and 4) an inversion on chromosome 14 (Figure 1A) ^20,21,28^. The asymmetric structure and chromosome-unique features around the SST1 arrays suggested we would be able to map the translocation site (Figure 1B). To further test this working model, we sequenced and assembled the complete, telomere-to-telomere sequence of three Robertsonian chromosomes.

**Figure 1.**
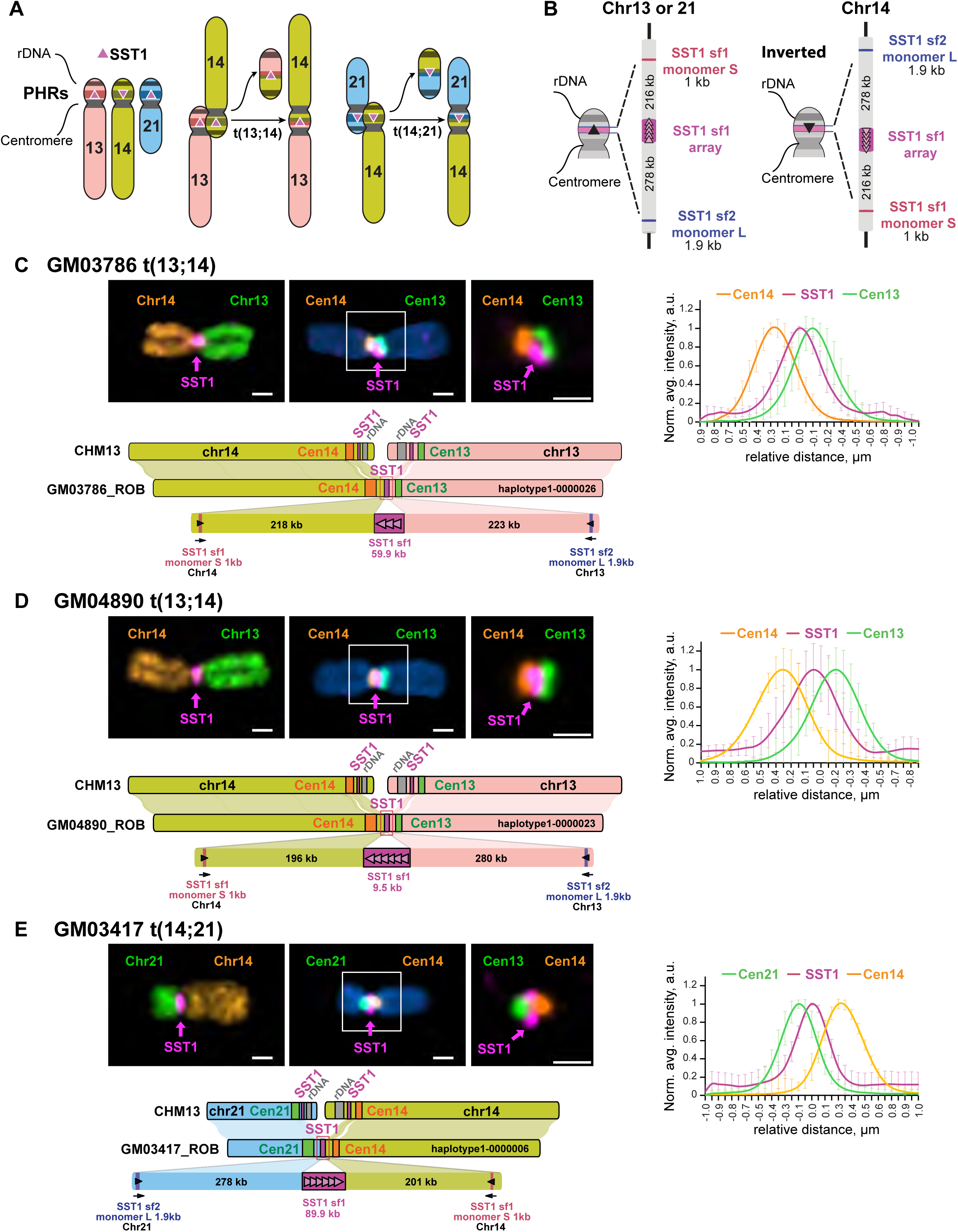
Complete assembly of ROBs. **A)** Working model for ROB formation depending on recombination between SST1 repeats (pink triangles) located in PHRs (colored bands) on the short arms of human chromosomes 13, 14, and 21. The adjacent 45S rDNA arrays, also called NORs, facilitate 3D proximity by co-locating in nucleoli. **B)** Schematic representation of the main SST1 arrays and flanking sequences in acrocentric chromosomes from CHM13. This region is similar on chromosomes 13 and 21, but inverted on chromosome 14. **C-E)** Representative images of ROBs from each cell line are shown. The left top panels display chromosomes labeled with an SST1 probe (magenta) and whole chromosome paints as indicated. The middle panels show chromosomes labeled with an SST1 probe (magenta) and centromeric satellite probes for CenSat 14/22 (orange) and CenSat 13/21 (green). DNA was counterstained with DAPI. The magnified inset (right) demonstrates SST1 localization between the two centromere arrays. Scale bar is 1 µm. The plot on the right shows averaged intensity profiles of lines drawn through the centromeres of at least 10 ROBs. Intensity profiles were aligned to the peak of the Gaussian of the SST1 signal and normalized to the maximum intensity of each channel. Error bars denote standard deviations. The lower panels show a synteny plots comparing the assembled ROB to CHM13. The structure of each fused region is shown in detail.

We selected three independent cell lines for sequencing and assembly, each harboring a unique ROB: 1) GM03417 (45,XX,der(14;21), clinical features of Down syndrome), 2) GM04890 (45,XX, dic(13;14)), clinically normal with 5 miscarriages), and 3) GM03786 (45,XX,dic(13;14), clinically normal). The fusions were initially confirmed in mitotic spreads using chromosome-specific paints (Figure 1C-E). Oxford Nanopore Technologies (ONT), Hi-C, ILMN, and PacBio HiFi sequencing data were collected. The Verkko assembler utilized ONT, PacBio, and Hi-C data to generate complete de novo assemblies of the ROBs^29^. Each ROB was visible in the assembly graph by a node connecting regions representing two different acrocentric chromosomes (Supplementary Figures 1-3). Notably, these connecting nodes skipped the rDNA arrays, consistent with the working model and supporting the loss of rDNA in the common ROBs.

We assessed the quality of the assemblies (Methods), finding high Phred quality scores (QVs) ranging from 49.10 to 54.70, corresponding to approximately 99.999% accuracy, and high gene completeness, with all assemblies containing over 98% of Benchmarking Universal Single-Copy Orthologs ^30^(BUSCOs) (Supplementary Figure 4). Read coverage analysis across the breakpoint regions shows consistent coverage without significant drops or high-frequency secondary alleles, confirming the structural accuracy of the assembled fusion points (Supplementary Figure 5). Hi-C data, which captures the 3-dimensional organization of chromosomes, showed increased interactions between the q arms of each ROB, emphasizing the physical proximity of these regions due to the fusion (Supplementary Figure 6).

SST1 macrosatellite arrays are present on several chromosomes ^31^. Multiple subfamilies of SST1 arrays exist. One subfamily occurs in the PHRs of chromosomes 13, 14, and 21, whose population genetic and structural features have led to the hypothesis that they are the site of recombination in recurrent Robertsonian fusions ^20^. Using cytogenetics, we probed centromeres 13/21, 14/22, and both SST1 and 45S rDNA arrays. Centromeres 13/21 and 14/22 are highly similar at the sequence level and cannot be distinguished by fluorescence in situ hybridization (FISH) probes ^32^. Our analysis confirmed that all three ROBs are missing 45S rDNA arrays and the SST1 signal is present between centromeric arrays (Supplementary Figure 7), consistent with the working model for the formation of common ROB and the assembly graphs.

Notably, the normal copy of chromosome 14 in GM03786 has lost the SST1 array (Supplementary Figures 2 and 7), suggesting polymorphisms in this region exist.

To visualize the assemblies, we created synteny plots (Figure 1C-E) using NGenomeSyn ^33^. SST1 arrays are present on 13, 14, and 21, but are inverted on chromosome 14 relative to chromosomes 13 and 21 in many genomes ^20^. Due to this inversion, when this region of chromosome 14 pairs with a 13 or 21 partner chromosome in meiosis, a crossover event is predicted to join the two long arms, forming a ROB. When we compared the assembled ROBs to their component chromosomes in CHM13, we observed that the sequence order and directionality was syntenic on either side of the SST1 region. This pattern is consistent with crossover within the SST1 array generating each of the assembled ROBs. Sequences unique to either chromosome 13/21 or 14, which include a partial SST1 monomer, allowed us to identify that the SST1 array was the breakpoint (Supplementary Figure 8).

### SST1 and evidence for interchromosomal exchange

To better understand the evolutionary relationships and potential exchange between SST1 arrays, we conducted a phylogenetic analysis of SST1 monomers derived from HG002 and CHM13 genomes. The analysis revealed three distinct subfamilies. Subfamily 1 (sf1), also known as NBL2, consists of monomers primarily derived from large arrays on the acrocentric chromosomes and form a single branch in the phylogenetic tree. Subfamily 2 (sf2) comprises monomers mainly from autosomal arrays (e.g., chromosomes 4, 17, and 19) and forms chromosome-specific branches. Subfamily 3 (sf3), also referred to as TTY2, is composed of monomers primarily originating from arrays on the Y chromosome. This classification of subfamilies and their distinct characteristics are illustrated in Figure 2A, 2C, and Supplementary Figure 8. The Y-derived repeats co-cluster with a few SST1 monomers from autosomes (see more below). The co-clustering of sf1 monomers from the large arrays on the acrocentric chromosomes is consistent with frequent recombinational exchange between these repeats, leading to concerted evolution of their constituent monomers.

**Figure 2.**
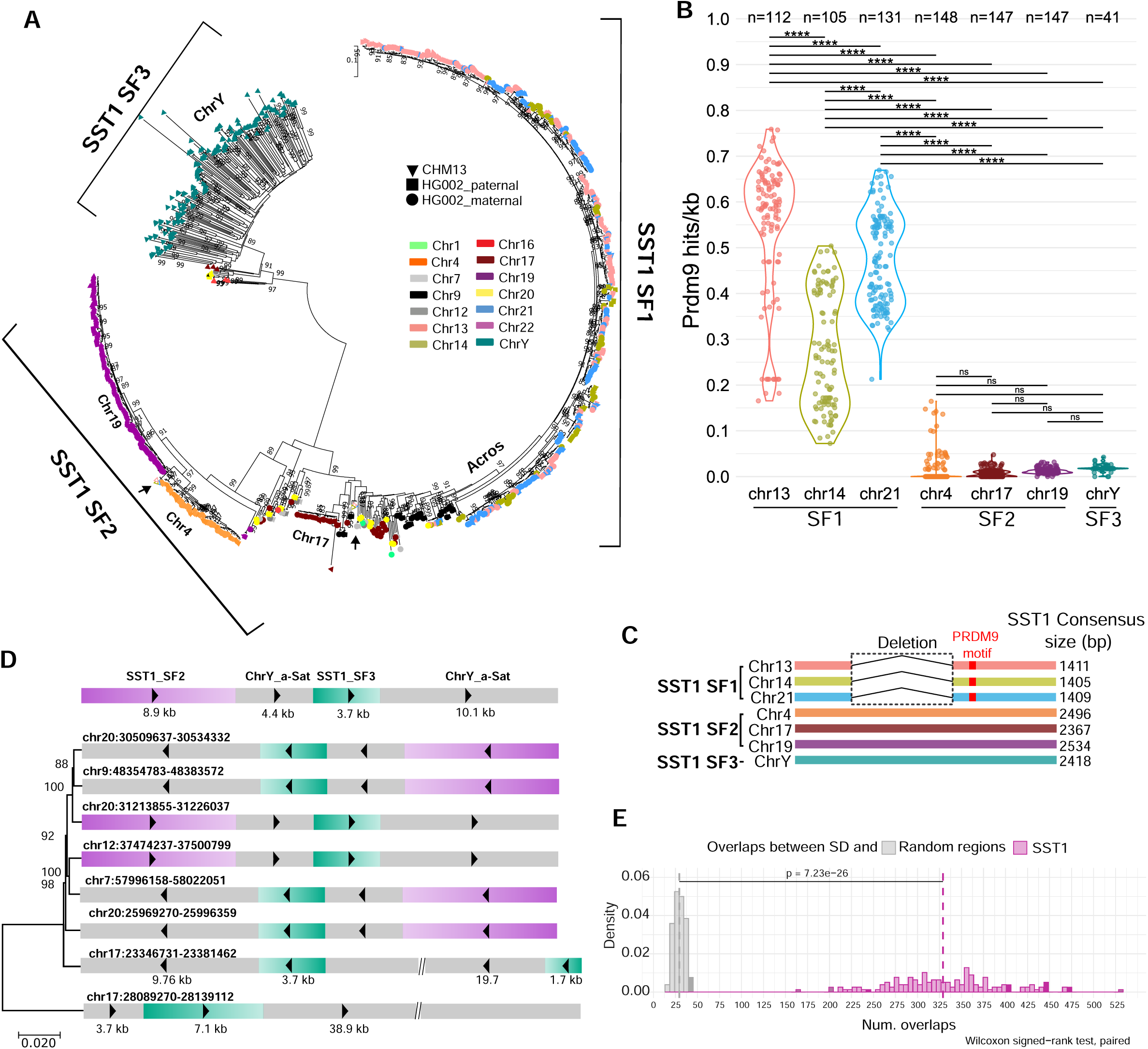
Evidence for SST1-mediated interchromosomal exchange in human genomes. **A)** All SST1 monomers from CHM13 and HG002 were collected and phylogenetic analysis was performed using the maximum likelihood method based on the best-fit substitution model (Kimura 2-parameter +G, parameter = 5.5047) inferred by Jmodeltest2 with 1,000 bootstrap replicates. Bootstrap values higher than 75 are indicated at the base of each node. The color indicates the source chromosome and the shape indicates the source genome. Three major subfamilies were identified: 1) subfamily 1, primarily on the acrocentrics, 2) subfamily 2, primarily on the remaining autosomes, and 3) subfamily 3, primarily on the Y chromosome. Black arrows indicate the location on the phylogenetic trees of sf2 monomers S and L from the acrocentric chromosomes (Figure 1B). **B)** Predicted PRDM9 DNA binding site frequency (mean sites/kb, each dot indicates 1 haplotype) in SST1 arrays in multiple haploid genomes (indicated by n), plotted by chromosome. ANOVA analysis with the Tukey-Kramer test for pairwise mean comparisons was used. **** indicates p<0.0001 and ns indicates not significant. **C)** Schematic representation of the three subfamilies of SST1. SST1 sf1 has a central gap and a predicted PRDM9 DNA binding site (red box). **D)** A segmental duplication of 27 kb or larger was identified on several autosomes in CHM13 that includes Y-like alpha-satellite DNA and Y-like SST1. Phylogenetic analysis was performed using the maximum likelihood method and GTR+Gamma substitution parameters. Bootstrap values are shown. **E)** Comparison of overlaps between segmental duplications (SD) and random regions (gray) or SST1 monomers (pink) across 147 genomes. Distributions show the number of overlaps (x-axis) versus density (y-axis). A permutation test with 10000 iterations per genome was used to generate random region overlaps. The significant difference between distributions (p value = 7.23e-26, Wilcoxon signed-rank test, paired) indicates non-random association between SD and SST1 regions.

The recurrent involvement of the SST1 array in ROB formation shown here and the concerted evolution of SST1 arrays on chromosomes 13, 14, and 21 ^20^ suggest that this array may comprise a meiotic recombination hotspot. To investigate this, we focused on PRDM9, a key protein in meiosis that plays a crucial role in defining recombination hotspots. PRDM9 contains a variable number of zinc fingers that allow it to bind to specific DNA sequences. Additionally, it possesses a histone methyltransferase domain that trimethylates histone H3 at lysine 4 (H3K4) and lysine 36 (H3K36) ^34,35^. This trimethylation of H3K4 and H3K36 by PRDM9 creates an open chromatin environment that favors recombination events during meiosis ^25^. Previous work suggested SST1 arrays on chromosomes 13, 14, and 21 in CHM13 contain predicted PRDM9 binding sites ^20^. To further examine PRDM9 binding sites within SST1 arrays, we searched for the 13-bp PRDM9 DNA binding motif ^36^ of the common A allele^37^ in the SST1 arrays in 147 human genomes (Methods). The density of PRDM9 DNA binding motifs is significantly higher in the SST1 arrays on chromosome 13, 14, and 21 relative to chromosomes 4, 17, 18, and Y (Figure 2B). Interestingly, chromosome 14 has a lower density compared to 13 and 21.

PRDM9 activity leads to the erosion of its own binding sites ^25^. PRDM9 sites in SST1 arrays should erode on all three acrocentric chromosomes. We speculate that PRDM9 sites within the SST1 arrays can regenerate via intra- or interchromosomal exchange events. However, the inversion on chromosome 14 may create a barrier for interchromosomal gene conversion, tipping the relative balance of erosion and regeneration toward erosion. The significantly higher density of PRDM9 DNA binding motifs in SST1 arrays on acrocentric chromosomes aligns with our finding that the consensus sequence of their monomers contains a predicted PRDM9 DNA binding motif (Figure 2C, Supplementary Figure 9). These findings suggest the predominant subfamily of SST1 repeats on acrocentric chromosomes (sf1) creates a sequence context permissive to meiotic recombination.

69 PRDM9 alleles have been documented in the human population ^37^. Some differ by a single nucleotide; others vary substantially by copy number and organization of their zinc-finger motifs. PRDM9 alleles can influence the activity of meiotic recombination hotspots ^38^. We reasoned that PRDM9 allele variation could influence the likelihood of ROB formation, so we examined the genotype of our three ROB cell lines. GM3417 is A/A, GM3786 is A/L24 and GM4890 is L24/L9. A is the most common allele but L24 and L9 are rare, present in only 1.66% of individuals in the previous study (12/720), although these alleles are overrepresented in an ethnically southern European population (9/109)^37^. In vitro assays show that L24 and L9 have a reduced binding affinity to many A-type binding motifs^39^ and can impact recombination activity at the MSTM1a hotspot ^38^. Any conclusions would be premature since we lack information on ancestry and vertical transmission in our samples.

Further evidence for SST1-mediated interchromosomal recombination comes from two key observations. First, the phylogenetic analysis revealed the presence of Y-like SST1 monomers (sf3) on multiple autosomes (7, 9, 12, 17, 20). These monomers are contained within larger sequence blocks ranging in size from 25-50 kb with 95-99% identity that also include Y-like alpha-satellite DNA (Figure 2D). Second, we examined the association between SST1 arrays and segmentally duplicated regions, defined as regions longer than 1 kb with at least 90% identity at two or more locations in the genome ^40^. Segmental duplications arise and persist due to recombination. On CHM13, we found complex patterns of homology, particularly between chromosomes 7, 9, 17, and 20. To quantify this association more broadly, we conducted a permutation test with 10000 iterations in 147 genomes (Methods). The results suggest the observed association between SST1 and segmental duplications is unlikely to occur by random chance (p value = 7.23e-26, Wilcoxon signed-rank test, paired) (Figure 2E). This pattern suggests these blocks have been copied from the Y chromosome, although we could not identify syntenic blocks on the human Y chromosome. Taken together, these findings suggest that, beyond its role in recurrent Robertsonian translocations, SST1 is associated with interchromosomal recombination throughout the genome.

### The association of SST1 arrays and NORs in chimp and bonobo genomes

Tandem arrays of the SST1 macrosatellite are not unique to humans; they also exist in nonhuman primate genomes, including several recently assembled telomere-to-telomere (T2T) genomes^41^. In gibbon, SST1 arrays have been suggested to be responsible for evolutionary patterns of genome instability^42^. As chimpanzees (*P. troglodytes*) and bonobos (*P. paniscus*) are the most closely related species to humans, we examined the acrocentric chromosomes in their genomes for the position and orientation of NORs and SST1 arrays to determine the incidence of the structural arrangement that gives rise to the common ROBs in humans. Two out of five NOR-bearing acrocentric p arms in the bonobo genome (hsa14 and hsa22) and all five NOR-bearing acrocentric p arms in the chimpanzee genome (hsa13, hsa14, hsa18, hsa21, hsa22) have co-oriented SST1 arrays (Figure 3A). The location of SST1 arrays on these chromosomes was validated by FISH (Supplementary Figure 10).

**Figure 3.**
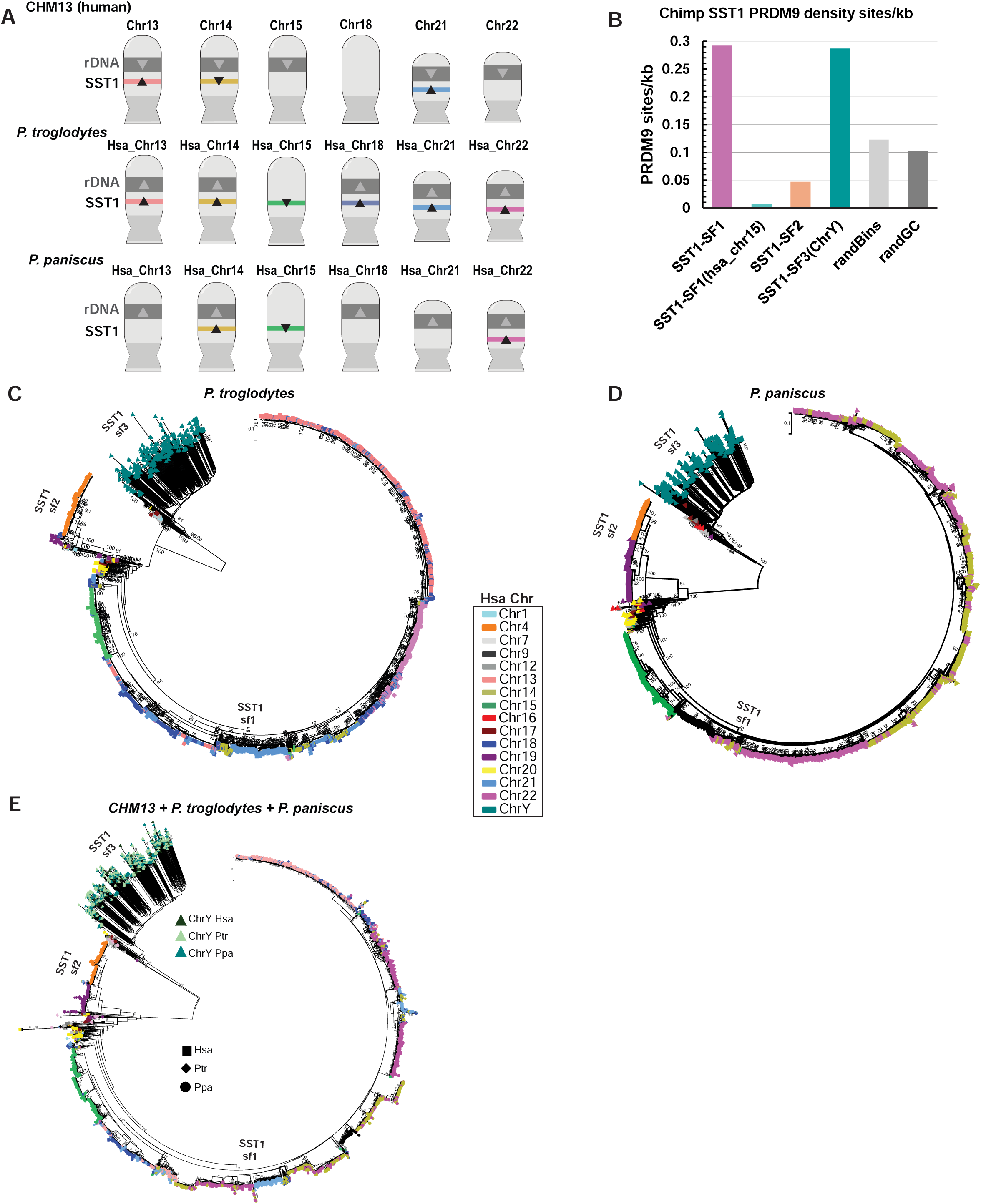
Evidence for exchange of SST1 on NOR-bearing chromosomes in chimpanzee and bonobo genomes. **A)** Ideograms of all the NOR-bearing chromosomes in human, chimpanzee, and bonobo, annotated with the human numbering system. The directionality of 45S rRNA gene arrays (grey) and SST1 arrays (colored bars) are indicated with arrowheads. **B)** Predicted PRDM9 binding sites were identified in the chimpanzee genome, and the number of sites per kb is plotted for SST1 arrays for the subfamily indicated. Random regions of the genome (randBins, randGC which was GC matched) were used to determine background. **C)** All SST1 monomers from the chimpanzee genome were subjected to phylogenetic analysis using the maximum likelihood method. The color indicates the source chromosome. Notably, the SST1 monomers from chromosomes 13, 14, 18, 21, and 22 form a single branch, indicating a high degree of similarity. **D)** All SST1 monomers from the bonobo genome were subjected to phylogenetic analysis using the maximum likelihood method. Notably, the SST1 monomers from chromosomes 14 and 22 form a single branch, indicating a high degree of similarity. **E)** SST1 monomers from human, chimpanzee and bonobo were subjected to phylogenetic analysis using the maximum likelihood method. The three subfamilies are apparent.

To investigate the potential role of PRDM9 in SST1-mediated recombination, we examined the frequency of predicted PRDM9 DNA binding sites^43,44^ in SST1 repeats in the chimpanzee genome. The density is high on the NOR-bearing chromosomes, but lower on the other chromosomes (Figure 3B). Interestingly, although the SST1 arrays on hsa15 in the chimpanzee genome are composed of subfamily 1 monomers, these monomers are not enriched for predicted PRDM9 DNA binding sites. This is consistent with the monophyletic state of SST1 from hsa15 in chimpanzees and bonobos (Figure 3C and 3D), which indicates that these arrays do not recombine with those on other NOR-bearing chromosomes. Altogether, our findings suggest that SST1 arrays can instigate exchange between nonhomologous chromosomes, especially in combination with NORs, shaping the evolution of primate genomes.

To gain insights into the evolutionary history of SST1 arrays and their potential role in chromosomal rearrangements across closely related species, we conducted a comparative phylogenetic analysis of SST1 monomers from NOR-bearing chromosomes in chimpanzee and bonobo. The analysis revealed patterns similar to those observed in the human genome. In the chimpanzee genome, monomers from hsa13, hsa14, hsa18, hsa21, and hsa22, co-cluster, while in the bonobo genome, monomers from hsa14 and hsa22 co-cluster (Figure 3C and 3D). These findings provide evidence for inter-chromosomal exchange between SST1 monomers on NOR-bearing chromosomes in both chimpanzee and bonobo genomes, suggesting that this phenomenon is not unique to humans but is a shared feature among great apes. When we performed a combined phylogenetic analysis of monomers from chimpanzee, human, and bonobo, we observed that the acrocentric chromosomes from all three species contain SST1 monomers of subfamily 1 (Figure 3E). The subfamily 3 monomers from the Y also co-cluster. Monomers from other parts of the genome form a separate branch for subfamily 2.

The inverted orientation of the SST1 array on the p arm of chromosome hsa15 in both chimpanzee and bonobo genomes is similar to the configuration of the SST1 array on chromosome 14 in the human genome (Figure 3A). However, the SST1 monomers from hsa15 form a chromosome-specific branch in the phylogenetic trees (Figure 3C-3E), indicating a lack of recombination between SST1 monomers on hsa15 and other chromosomes in the ape genomes. This observation may be explained by the absence of a NOR on hsa15 in the chimpanzee and bonobo, which we speculate brings the SST1-containing regions in proximity to the nucleolus, facilitating interchromosomal exchange. Consequently, the chimpanzee and bonobo genomes lack the specific placement and orientation of NORs and SST1 arrays that we hypothesize facilitate the formation of human common ROBs, suggesting that this structural arrangement is unique to the human genome. Correspondingly, the presence of the NOR and absence of SST1 on human chromosome 15 suggests that the ancestral hsa15 of human, chimpanzee, and bonobo may have had both NOR and SST1 arrays. The greater sequence divergence of chromosome 15 from other NOR-bearing chromosomes in humans highlights the importance of SST1 for maintaining their sequence similarity^20^.

The association between SST1 and interchromosomal recombination events is further supported by the identification of regions syntenic with the segmental duplication shown in Figure 2D across several nonhuman primate genomes (Supplementary Figure 11A). This finding suggests that these duplication events occurred in a common ancestor. Moreover, we identified a second segmental duplication on chromosome 16 in primates (hsa16), which exhibits a different structure but also contains Y-like SST1 and alpha satellite sequences (Supplementary Figure 11B). These findings provide compelling evidence that SST1 is associated with interchromosomal recombination in primate genomes, and past events have involved the Y chromosome as a donor. Notably, the ancestral version of the Y chromosome was likely NOR-bearing^45^.

### ROBs can be functionally mono- or di-centric

All ROBs examined in this study have two centromeric DNA arrays, roughly 5 Mb apart, yet they are faithfully transmitted through mitosis. This observation, consistent with previous cytogenetic observations^27,46^, suggests that epigenetic alterations may occur to the centromeres to enable correct chromosome transmission. To investigate centromere activity in ROBs, we leveraged both microscopy and genomic data. To analyze ROBs by microscopy, we performed immunoFISH using FISH probes specific to cen13/21 and cen14/22, along with antibodies against CENP-C, an inner kinetochore protein that marks the active centromere^47^, and CENP-B, which binds to the 17 bp CENP-B box in alpha satellite DNA^48^. CENP-B was present at all centromeres, consistent with previous reports for a dicentric chromosome^49^. Confocal fluorescence microscopy (Supplementary Figure 12) revealed that the CENP-B signal was proportional to the size of the centromeric array determined by the assembly (Supplementary Figures 13).

To gain more detailed insights into centromere activity in ROBs, we used super-resolution microscopy (SIM) to evaluate the localization of CENP-C in the two 13;14 fusion chromosomes. CENP-C signal co-localized with cen14 (Figure 4A-B), suggesting that cen14 is the active centromere, while cen13 is inactive. This observation is consistent with a previous study that found cen14 to be the active centromeric array in most 13;14 dicentrics^27^. The active array did not depend strictly on size, as in GM04890 cen14 was smaller than cen13 (Supplementary Figure 13). In contrast, the CENP-C localization pattern for the 14;21 fusion was less binary. CENP-C appeared to colocalize with both cen14 and cen21 within a single chromosome (Figure 4C), and the pattern exhibited some heterogeneity between chromosomes, suggesting both arrays may remain active.

**Figure 4.**
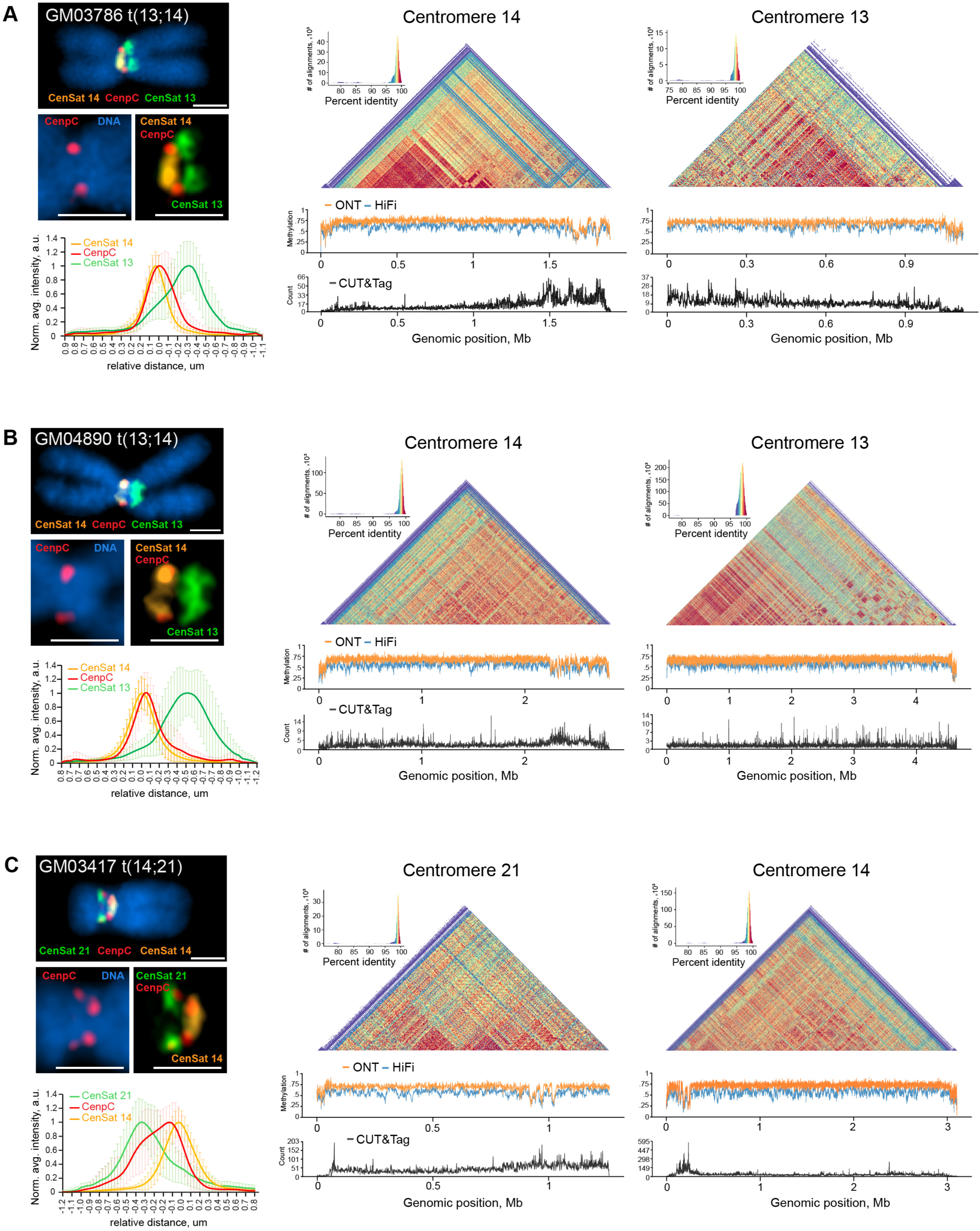
Centromere activity in dicentric ROBs. ImmunoFISH, DNA methylation, and CENP-A CUT&Tag analyses were performed for **(A)** GM03486, **(B)** GM04890 and **(C)** GM03417. The left panels show representative structured illumination super-resolution images of ROBs labeled by immuno-FISH with centromeric satellite probes for CenSat 14/22 (orange), CenSat 13/21 (green), and anti-CENP-C antibody (red). DNA was counterstained with DAPI. Magnified insets depict single CENP-C foci on centromere 14 in GM03786 and GM04890, and double CENP-C foci on centromeres 21 and 14 in GM03417. Scale bar is 1 µm. Plots below show averaged intensity profiles of lines drawn through the centromeric regions of sister chromatids of at least 10 ROBs. Intensity profiles were aligned to the peak of the Gaussian of the CenSat 14 signal and normalized to the maximum intensity of each channel. Error bars denote standard deviations. The right panels display corresponding heatmaps of sequence similarity calculated for 5 kb bins for each centromere. Below the heatmaps, DNA methylation tracks show methylation calls from ONT (orange) or PacBio HiFi (blue) sequencing, with hypomethylated regions suggesting active centromere localization. Active centromere regions are confirmed by CENP-A peaks on CUT&Tag tracks below (black).

To further investigate centromere activity in ROBs, we examined epigenetic markers associated with active centromeres. Previous studies have identified a dip in CpG methylation in active centromere arrays that coincides with enrichment or CENP-A, the centromere histone H3 variant. This region is referred to as the centromere dip region (CDR) and likely marks the site of kinetochore assembly^50,51^. We inspected CpG methylation from both HiFi and ONT reads across the centromere arrays in the three assembled ROBs. As controls, we examined CDRs in chromosomes 13, 14, and 21 from the normal chromosomes in the same cell line. We visualized CDRs in conjunction with pairwise sequence similarity heatmaps between 5 kb bins of the centromeric array^52^ and both methylation and CUT&Tag data for CENP-A. In the 13;14 fusions, there is a CDR on chromosome 14, coincident with CENP-A enrichment, and while there is low CENP-A signal in the adjacent cen13 array, there is no corresponding CDR (Figures 4A-B). This finding is consistent with the localization of CENP-C with the cen14 FISH probe, suggesting that cen14 is the active centromere and this chromosome is functionally monocentric. In contrast, cen13 on the normal chromosome has a clear CDR and CENP-A signal (Supplementary Figure 14).

In the 14;21 fusion, we observed a dip in methylation in both cen14 and cen21 arrays, located near the adjacent edges of the two arrays (Figure 4C) towards the fusion breakpoint. The CENP-A signal is stronger on cen14, but CENP-A signal also exists on cen21 coincident with a CDR. In combination with imaging, the data suggest this chromosome is functionally dicentric. The CDR and CENP-A signal on cen21 on the normal chromosome are more pronounced (Supplementary Figure 14). Previous studies have shown that functionally dicentric chromosomes with two active arrays can be stably transmitted through mitosis in the case of isodicentric X chromosomes, especially when centromeres are less than 12 Mb apart^26^. Our findings suggest that ROBs are stably propagated due to epigenetic adaptation in centromere activity. In some cases, this is achieved by complete inactivation of one array, while in other cases, activity is shared between two arrays.

## Discussion

While 15% of human ROBs appear to form by idiosyncratic mechanisms, 85% involve chromosome 14^17^. These common ROBs likely arise due to a unique combination of factors, including the inversion on chromosome 14, the presence of a NOR which draws the acrocentric short arms into physical proximity near nucleoli, and the recombination initiation from a hotspot at or near the SST1 array. Tandem arrays of SST1 sf1 are found primarily on NOR-bearing chromosomes in chimpanzee, bonobo, gorilla, and human genomes^41^. However, the assembled chimpanzee and bonobo genomes do not possess a chromosome containing both an inverted SST1 array and a NOR, suggesting that this particular structural arrangement is unique to the human lineage. Despite this, phylogenetic evidence suggests that interchromosomal exchange between SST1 arrays on NOR-containing chromosomes is a common feature in human, bonobo, and chimpanzee genomes. Furthermore, the SST1 sf1 arrays that show evidence of exchange in the human and chimpanzee genomes are also enriched for PRDM9 DNA binding sites.

Taken together, the evidence suggests that the common ROBs in humans occur via a combination of four key factors: 1) homologous SST1 repeats on nonhomologous chromosomes, 2) the inversion on chromosome 14, which creates a unique structural arrangement, 3) the co-location of two of these regions in 3D space due to the presence of NORs that bring the regions into close proximity (within the nucleolus), and 4) meiotic recombination hotspots, potentially mediated by PRDM9, that lead to crossing over between the nonhomologous chromosomes^21^.

Our work consolidates many disparate observations regarding the SST1 macrosatellite family ^22,23^, efforts that have been limited by the lack of repetitive DNAs in reference genomes. Through careful analysis of multiple human, chimpanzee, and bonobo genomes, we identified three distinct subfamilies of SST1. Several names appear in the literature, including TTY2^53^ or MER22 ^54^ for subfamily 3 on the Y chromosome, and NBL2^24,55–59^ for subfamily 1 on the acrocentric chromosomes. In several cases, SST1 sequence has been associated with genome instability and adverse health impacts. For example, loss of SST1 repeats on the Y is associated with male infertility^53^, while hypomethylation is associated with cancer^60,61^, potentially contributing to its transcription and subsequent genomic instability^62,63^. Our work is consistent with previous proposals that SST1 contributes to genome instability, and in fact, may have a much broader role than previously appreciated.

SST1 sequences on the acrocentric chromosomes are highly enriched for PRDM9 DNA binding sites, suggesting the creation of meiotic recombination hotspots that result in recombination between nonhomologous chromosomes and the formation of common ROBs. We speculate that PRDM9 DNA binding sites lost via erosion^25^ may be “regenerated” by gene conversion or other exchange events, allowing SST1-hotspot associated activity to persist and self-sustain. It is also possible that rare and functionally distinct PRDM9 alleles could contribute to the incidence of ROB formation and segmental duplications, based on their binding properties, but more studies will be required. Taken together, the evidence suggests SST1 repeats play a significant role in the stability and evolution of primate chromosomes, as first suggested for the gibbon genome^42^.

ROBs form more commonly in female meiosis^15,16^. Meiosis is sexually dimorphic in many ways that could contribute to the higher frequency of ROB formation in female meiosis as compared to male meiosis. Female meiosis proceeds through the earliest stages of prophase with DNA in a more demethylated state than in male meiosis^64^, and open chromatin is more prone to recombination. Consistently, there is more recombination initiation and crossing over in female meiosis^65–67^. Furthermore, hotspots can be sexually dimorphic^68^. The ends of chromosomes synapse later in female meiosis compared to male meiosis^66^, which could make them more prone to interchromosomal exchange. Future efforts will be required to distinguish between the transcriptional, epigenetic, and hotspot status of the short arms of acrocentric chromosomes in developmental and disease states in males and females, and the factors that allow ROBs to occur more commonly in female meiosis.

ROBs are transmitted at a higher rate than Mendelian ratios in female meiosis, a phenomenon known as meiotic drive^69^. Drive can occur in females because of the segregation of material into “dead end” polar bodies^70^, potentially due to weaker centromeres^71^. The novel structure of each ROB, including centromere activity, may impact its transmission. In our study, the 14;21 fusion chromosome appeared functionally dicentric, which may create a stronger centromere, potentially influencing its transmission. Moreover, ROBs are larger than each corresponding normal chromosome but have lost SST1-distal sequences. The number of crossovers on the ROB should be higher relative to the normal chromosome^72^, facilitating its segregation. However, the loss of SST1-distal sequences (i.e. rDNA arrays) may counteract this bias. Normal chromosomes will have unsynapsed regions in the trivalent synaptonemal complex structure containing the ROB^73,74^, which may trigger checkpoints^75^. The structure of each ROB will differ based on variation in individual centromeres and other repeats, potentially conferring different advantages and disadvantages for transmission. Analysis of many ROBs will be required to understand how individual features of ROBs impact their propagation and carrier fertility. By integrating insights from genomic, cytogenetic, and evolutionary studies, we can gain a more comprehensive understanding of the role of NORs and SST1-based recombination in human genome evolution and reproductive biology.

## Acknowledgements

For generating sequencing data, we thank Kate Hall, Michael Peterson, and the National Institutes of Health Intramural Sequencing Center (NISC). JLG, TPO, LGL, CS, SM are supported by the Stowers Institute for Medical Research. JLG, TPO and LGL are also supported by NCI R01CA266339. This work was supported, in part, by the Intramural Research Program of the National Human Genome Research Institute, US National Institutes of Health. This work utilized the computational resources of the NIH HPC Biowulf cluster (https://hpc.nih.gov). EG and AG are supported by NIH R01HG013017, NIH U01DA057530, NSF #2118743, and the State of Tennessee’s Center for Integrative and Translational Genomics. For helpful discussion, we thank Scott Hawley, Aurora Ruiz-Herrera, SaraH Zanders, Chuck Langley, and Terry Hassold.

## Data and Code Availability

All data derived from de-identified human patient DNA samples. Original raw data files for Sequencing are available via dbgap in accordance with the patient consent from Coriell. Cytogentic analyses are publicly available and can be accessed from the Stowers Original Data Repository at http://www.stowers.org/research/publications/libpb-XXXX

## Materials and methods

### Cell culture

Human lymphoblastoid cell lines (LCL) GM03786, GM04890, and GM03417 were obtained from Coriell. All LCL cell lines were cultured in RPMI 1640 (Gibco) with L-glutamine supplemented with 15% fetal bovine serum (FBS) in a 37°C incubator with 5% CO2.

| **Cell line** | **Platform** | **Sequencing depth** | **Accession Number** |
| --- | --- | --- | --- |
| **GM03786** | **PacBio Sequel II HiFi mode** | 53.74 | **TBD** |
|  | **ONT Nanopore** | 72.44 | **TBD** |
|  | **Illumina** | 55.89 | **TBD** |
|  | **Hi-C** | 57.88 | **TBD** |
| **GM04890** | **PacBio Sequel II HiFi mode** | 49.37 | **TBD** |
|  | **ONT Nanopore** | 75.24 | **TBD** |
|  | **Illumina** | 39.13 | **TBD** |
|  | **Hi-C** | 82.28 | **TBD** |
| **GM03417** | **PacBio Sequel II HiFi mode** | 56.23 | **TBD** |
|  | **ONT Nanopore** | 63.07 | **TBD** |
|  | **Illumina** | 51.27 | **TBD** |
|  | **Hi-C** | 65.29 | **TBD** |

### ONT sequencing

The UHMW DNA was extracted from frozen cell pellets using the NEB Monarch HMW DNA Extraction Kit for Tissue and assessed for fragment size using a pulsed field gel. The fragments span from 50 kb to 1000 kb. Genomic DNA libraries were prepared using the “NEB_5ml_Ultra-Long Sequencing Kit (SQK-ULK001)-promethion” protocol from Oxford Nanopore. Each library was loaded onto a FLO-PRO002 flow cell and ran for 72 hours with two subsequent loadings at 24-hour intervals. The libraries were sequenced on a PromethION (Oxford Nanopore) running MinKNOW software version 22.12.5. Basecalling and modified base detection (5mC) were performed on-instrument using Guppy 6.4.6 with the following model: dna_r9.4.1_450bps_modbases_5mc_cg_hac_prom.cfg..

### PacBio HiFi sequencing

PacBio library preparation was conducted using the SMRTBELL Prep Kit 3.0. The prepared libraries were quantified and sequenced on a PacBio Revio system with Instrument Control Software version 12.0.0.183503 and chemistry version 12.0.0.172289. Sequencing was performed using two SMRT Cells, each with a movie length of 24 hours.

Using Pacific Biosciences SMRTbell® Prep Kit 3.0 with binding kit 102-739-100 and sequencing kit 102-118-800, three libraries (one per sample). The Megarupter (Diagenode) was used for shearing and SageELF (Sage Science) was used for size-selection. Library size was assessed using a FemtoPulse (Agilent). Each library was run on version 25M SMRTCells using the first generation polymerase and chemistry version 1 (P1-C1). Sequencing was performed on a PacBio Revio system running instrument control software version 12.0.0.183503 and a movie collection time of 24 hours per SMRTCell. Using PacBio SMRTLink version 12.0.0.172289, CCS/HiFi reads generated on-instrument using ccs v7.0.0, lima v2.7.1 (demultiplexing), and primrose v1.4.0 (5mC calling).

### Hi-C sequencing

HiC libraries were generated according to manufacturer’s directions using the Arima High Coverage HiC User Guide for Mammalian Cell Lines (Document Part No. A160161 v01) and Arima-HiC+ User Guide for Library Preparation Using the Arima Library Prep Module (Document Part No. A160432 v01). Starting with 5M cells per sample, the Standard Input Crosslinking protocol was followed resulting in 1.49-1.86ug of DNA available per sample to generate large proximally-ligated DNA as assessed using the Qubit Fluorometer (Life Technologies). Library preparation was performed using the S220 Focused-ultrasonicator (Covaris) to shear samples to 550bp followed by a DNA purification bead cleanup with no size selection, and 5 or 7 cycles of library PCR amplification per sample. Resulting short fragment libraries were checked for quality and quantity using the Bioanalyzer (Agilent) and Qubit Fluorometer (Life Technologies). Libraries were pooled, requantified and sequenced as 150bp paired reads on both the Illumina NextSeq 2000 and NextSeq 500 instruments to obtain at least 600M read pairs per sample, utilizing RTA and instrument software versions current at the time of processing. Demultiplexing was performed with bcl-convert v3.10.5. The cut sites for the enzymes used were ^GATC, G^ANTC, C^TNAG, and T^TAA.

### ILMN (GM04890, GM03786)

Libraries were generated from 100 nanograms of genomic DNA using Covaris LE220 plus to shear the DNA and the 2S™ Plus DNA Library Kit (Integrated DNA technologies cat.# 10009878) for library preparation. To minimize coverage bias only 4 cycles of PCR amplification were used. The median insert sizes were approximately 300 bp. Libraries were tagged with unique dual index DNA barcodes to allow pooling of libraries and minimize the impact of barcode hopping. Libraries were pooled for sequencing on the NovaSeq X plus (Illumina) across 14 lanes to obtain at least 369 million 151-base read pairs per individual library.

### ILMN (GM03417)

PCR-free libraries were generated from 1 microgram genomic DNA using a Covaris R230 to shear the DNA and the TruSeq® DNA PCR-Free HT Sample Preparation Kit (Illumina) for library preparation. The median insert sizes were approximately 400 bp. Libraries were tagged with unique dual index DNA barcodes to allow pooling of libraries and minimize the impact of barcode hopping. Libraries were pooled for sequencing on the NovaSeq X plus (Illumina) across 7 lanes on 25B flowcells to obtain at least 388 million 151-base read pairs per individual library.

### Assembly methods

Phased genome assemblies were generated using Verkko (v1.4.1)^1^. The assembly process integrated PacBio HiFi reads and Oxford Nanopore (ONT) reads, with Hi-C reads used specifically for the phasing. The ONT reads included ultra-long reads, defined as reads that are at least 100 kb in length. Verkko was run with the command:

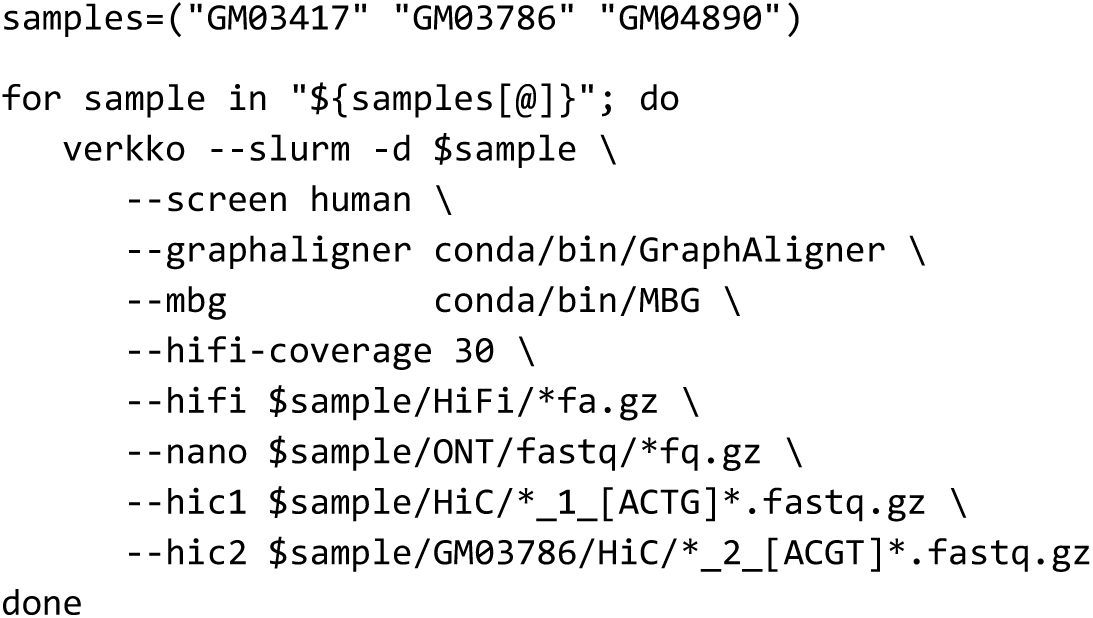

Haplotype-consistent contigs and scaffolds were automatically extracted from the labeled Verkko graph, with unresolved gap sizes estimated directly from the graph structure. After the assembly was generated, we collapsed all nodes composed of only rDNA k-mers into a single node and added telomere nodes to the graph to indicate ends of chromosomes using the commands:

~~~
seqtk hpc rDNA.fasta > rDNA_compressed.fasta
seqtk telo assembly.fasta > assembly.telomere.bed
mash sketch -i 8-hicPipeline/unitigs.hpc.fasta -o compressed.sketch.msh
$mash screen compressed.sketch.msh rDNA_compressed.fasta | awk ’{if ($1 > 0.9 && $4 < 0.05) print $NF}’ >
target.screennodes.out
python remove_nodes_add_telomere.py -r target.screennodes.out -t assembly.telomere.bed
~~~

In this simplified graph, the Robertsonian translocation was apparent in all cases (Supplementary Figures 1-3). We extracted the assembly path corresponding to the Robertsonian chromosome and identified gaps in the assembly. There was 1 gap in GM03417, 1 gap in GM03786, and 2 gaps in GM04890. Manual interventions were employed to complete the chromosomes.

### Assembly quality evaluation

We evaluated the quality and gene completeness of the genome assemblies using two approaches: a k-mer-based, reference-free method and a gene content assessment. For the k-mer-based evaluation, we employed Merqury^2^, a tool that assesses assembly completeness and accuracy without relying on a reference genome. Merqury uses k-mer frequencies from sequencing reads to estimate the quality value (QV) of the assemblies, which represents the phred-scaled error rate. For our evaluation, we used PacBio HiFi reads for the QV estimation.

To assess gene completeness, we used compleasm^3^, a tool based on BUSCO (Benchmarking Universal Single-Copy Orthologs). Compleasm evaluates the presence and integrity of a curated set of BUSCOs expected to be present in the genomes of the taxonomic group under study. We used the primate-specific BUSCO dataset, which includes 13780 genes, to quantify the completeness, duplication, and fragmentation of conserved genes in our assemblies.

### PRDM9 site predictions and density

In 147 human haploid genomes (from 72 diploid individuals plus the haploid CHM13 and the diploid HG002), predicted PRDM9 DNA binding sites were identified by using Motifence (v0.1.1, commit fb1ebc0, https://github.com/AndreaGuarracino/motifence) to find DNA sequences matching the canonical 13-mer motif CCNCCNTNNCCNC^4^ or its reverse complement. To compute the density of PRDM9 DNA binding sites per kilobase (kb) in SST1 regions, SST1 arrays were first identified using TideHunter^5^. For a region to be defined as an SST1 array, the following criteria were applied: monomeric unit within the array had to be at least 500 base pairs (bp) in length, there had to be at least two monomers, and the monomers had to overlap with RepeatMasker (v4.1.5, http://repeatmasker.org/) SST1 annotations. The PRDM9 density was then calculated by dividing the number of PRDM9 binding sites in the SST1 regions by the total length of these SST1 regions. PRDM9 alleles were found by conducting a BLAST search (blast-plus/2.13.0) on GM3417, GM3786 and GM4890 with the A allele as the reference. To identify genotypes, these hits were aligned to the 69 alleles from Alleva et al^6^ using MUSCLE and visualized in Geneious Prime 2024.0.7.

In the chimpanzee genome, PRDM9 site density in sites per kb on SST1 regions was calculated using R and Bioconductor. The function vmatchPattern from the Biostrings library was used to map the occurrence of the chimpanzee PRDM9 motifs: prdm9_E CNNCCNAANAA, prdm9_W CNGNNAANANTT, prdm9_pt1 ANTTNNATCNTCC, or their reverse compliments, on the genome. SST1 containing regions were then queried for overlap of PRDM9 sites using the countOverlaps function from the GenomicRanges library. Query width was used to calculate sites/kb. SST1 regions larger than 10kb were broken into 3 kb tiles to approximate resolution near SST1 feature size. Background PRDM9 site density was assessed in two ways. Random background PRDM9 density for each chromosome was determined using 100 randomly chosen 3kb segments. To account for GC bias, the genome was scored for GC content at 3 kb resolution, and fragments within one standard deviation of the average GC content of the SST1 containing elements were chosen to calculate background PRDM9 site density.

### SST1-Segmental Duplication association

To examine associations between SST1 repeats and segmental duplications, we performed the following analysis in 147 human genomes (from 72 diploid individuals plus the haploid CHM13 and the diploid HG002). First, repetitive regions in the genomic sequences were masked using RepeatMasker (v4.1.5, http://repeatmasker.org/) and Tandem Repeats Finder (v4.09.1)^7^. Segmental duplications were then identified using SEDEF (v1.1)^8^, on each haploid masked genome. SST1 repeats were detected using RepeatMasker and refined with TideHunter, as described above. Finally, we used the R package regioneR (v1.36.0)^9^, to perform permutation testing (n=10000) to assess the significance of spatial associations between SST1 repeats and segmental duplications. This analysis was conducted on 147 haplotype-resolved genomes to provide a comprehensive view of these genomic features across diverse human genomes.

### SST1 monomer characterization

We used RepeatMasker to find the regions. We retrieved all fasta files with 1kb of flanking regions for all arrays. Then, we manually curated all clusters using visual inspection by generating dot-plots with the Dotlet applet^10^ with a 15 bp word size and 60% similarity cut-off. We made regressive changes in the consensus sequences used and that enabled us to describe the sequences properly. By manual curation, we were able to identify the beginning and end of the arrays and each monomer regarding the consensus generated. All monomeric sequences analyzed were characterized with the same initial and final point regarding the consensus for the sake of alignment.

### Maximum likelihood phylogenetic analysis

We aligned all SST1 full-length monomeric sequences retrieved from assembled genomes using MUSCLE^11^. We conducted the phylogenetic analysis by using the maximum likelihood method based on the best-fit substitution model (Kimura 2-parameter +G, parameter = 5.5047) inferred by Jmodeltest2 with 1,000 bootstrap replicates. Bootstrap values higher than 75 are indicated at the base of each node.

### Chromosome spreads, Fluorescent In-Situ Hybridization (FISH), and immuno-FISH

For the preparation of chromosome spreads, cells were blocked in mitosis by the addition of Karyomax colcemid solution (0.1 µg/ml, Life Technologies) for 6-7h. Adherent fibroblast cells were collected by trypsinization. Collected cells were incubated in hypotonic 0.4% KCl solution for 12 min and prefixed by addition of methanol:acetic acid (3:1) fixative solution (1% total volume). Pre-fixed cells were spun down and then fixed in Methanol:Acetic acid (3:1).

For SST1 and CenSat FISH, spreads were dropped on a glass slide and incubated at 65°C overnight. Before hybridization, slides were treated with 0.1mg/ml RNAse A (Qiagen) in 2xSSC for 45 minutes at 37°C and dehydrated in a 70%, 80%, and 100% ethanol series for 2 minutes each. Slides were denatured in 70% deionized formamide/2X SSC solution pre-heated to 72°C for 1.5 min. Denaturation was stopped by immersing slides in 70%, 80%, and 100% ethanol series chilled to -20°C. Labeled DNA probes were denatured separately in a hybridization buffer by heating to 80°C for 10 minutes before applying to denatured slides. Fluorescently labeled human CenSat probes for D13Z1/D21Z1 and D14Z1/D22Z1 were from Cytocell. The biotin-labeled BAC probe for SST1 (RP11-614F17) was obtained from Empire genomics. Specimens were hybridized to the probes under a glass coverslip or HybriSlip hybridization cover (GRACE Biolabs) sealed with the rubber cement or Cytobond (SciGene) in a humidified chamber at 37°C for 48 -72hours. After hybridization, slides were washed in 50% formamide/2X SSC 3 times for 5 minutes per wash at 45°C, then in 1x SSC solution at 45°C for 5 minutes twice and at room temperature once. For biotin detection, slides were incubated with fluorescent streptavidin conjugated with Cy5 (ThermoFisher Scientific) for 2-3 hours in PBS containing 0.1% Triton X-100 and 5% bovine serum albumin (BSA), and then washed 3 times for 5 minutes with PBS/0.1% Triton X-100. Slides were mounted in Vectashield containing DAPI (Vector Laboratories). Confocal Z-stack images were acquired on the Nikon TiE microscope equipped with PlanApo 100x oil immersion objective NA 1.45, Yokogawa CSU-W1 spinning disk, Flash 4.0 sCMOS camera (Hamamatsu), and NIS Elements software.

For CENP-B and CENP-C immuno-FISH, freshly prepared chromosome spreads were dropped on a glass slide, washed with PBS/0.1% Triton X-100, and blocked with 5% bovine serum albumin (BSA) in PBS/0.1% Triton X-100. Primary antibody (rabbit polyclonal anti-CENP-B, Abcam, cat.# ab25734, rabbit polyclonal anti-CENP-C, Millipore, cat.# ABE1957) and secondary antibody (goat anti-rabbit Alexa Fluor 647, ThermoFisher Scientific) were diluted in 5% BSA/PBS/0.1% Triton X-100. Specimens were incubated with primary antibody overnight, washed 3 times for 5 minutes, incubated with secondary antibody for 2-4 hours and washed again 3 times for 5 minutes. All washes were performed with PBS/0.1% Triton X-100. After antibody incubation, spreads were post-fixed in 2% paraformaldehyde diluted in PBS for 15 min, washed in PBS, and processed for CenSat FISH as described above, starting with ethanol dehydration series. DNA was stained with 1.5 µg/ml DAPI. Confocal Z-stack images of CENP-B immuno-FISH were acquired on the Nikon TiE microscope as described above. For structured illumination super-resolution microscopy (SIM) performed on CENP-C immuno-FISH, slides were rinsed in ddH2O, air-dried in the dark, mounted in ProLong Glass antifade mountant (ThermoFisher Scientific) and allowed to cure for at least 24 hours before imaging. Z-stack images were acquired on an Elyra 7 Lattice SIM^2^ microscope (Zeiss) equipped with two PCO.edge 4.2 sCMOS cameras, four high power continuous wave lasers (405, 488, 561 and 642 nm), and a Zeiss PlanApo 63x oil immersion objective NA 1.4. The illumination pattern was set to 15 phases, and the Z-stack spacing was set at 100 nm. Raw SIM images were reconstructed using the ZEN Black software (Zeiss) with 10.5 manual adjustments for sharpness and best fit settings for all channels except 405nm (DAPI), which was processed in the widefield mode.

### Centromere intensity profiling of CenSat and SST1 FISH and CENP-B immuno-FISH

Maximum intensity projections from spinning disk confocal Z-stacks were generated, and chromosomes of interest were segmented manually based on DNA and CenSat labeling. Segmented chromosomes from each cell line were oriented vertically and assembled in a new stack consisting of identified specific chromosomes from multiple chromosome spreads. Intensity plot profiles were generated from 2µM vertical lines with the width of 10 pixels drawn through centromeric regions of each chromosome. Intensity profiles were combined by channel, fit to single Gaussian functions, and aligned to the peak of the Gaussian of the indicated channel. These profiles were then averaged together and normalized to the maximum intensity of each peak. For each chromosome from each cell line, at least 10 intensity profiles were averaged and plotted with standard deviations. All image processing and analysis were performed using ImageJ/FIJI. A detailed description of this type of analysis and relevant plugins are available at https://research.stowers.org/imagejplugins/spasim.html.

### Semi-automated intensity profiling of CENP-C immuno-FISH from SIM images

Reconstructed SIM images were mean projected, except for the DAPI channel, which had the slice of highest contrast selected. Robertsonian chromosomes and corresponding normal acrocentric chromosomes were identified using CenSat FISH signals and segmented manually or with a Cellpose model trained on a combination of the DAPI and CenSat signals. Individual chromosomes were transferred to a new image and oriented vertically using a second Cellpose model trained to find a skeleton of the chromosome. Bent chromosomes were straightened in ImageJ/FIJI by manually drawing two annotation lines across centromeres, one through each kinetochore. The straightened images were then aligned to the peak of the specified CenSat FISH signal used as the anchor point, and the line intensity profiles were aggregated over multiple images and split by cell line. At least 10 chromosomes were analyzed for each instance from each cell line. All analysis was performed in ImageJ/FIJI and python with code at the following: https://github.com/jouyun/Gerton_Robertsonian_2024.

### Methylation calls

HiFi BAM and ONT FASTQ files with 5mC methylation calls as MM and ML tags were aligned against the generated assemblies using pbmm2 (v.1.13.0, https://github.com/PacificBiosciences/pbmm2) for HiFi reads and Winnomap (v2.03)^12^, for ONT reads. The alignments were then converted to sorted BAM files containing only primary mappings with samtools (v1.17)^13^:

~~~
# HiFi reads
pbmm2 align {genome}.mmi {bam_with_meth_calls} -j 42 > {output.bam}
samtools view -@ 24 -Sb -F 2048 {output.bam} | samtools sort -@ 24 -T {temporary_directory} - > {output.bam}
samtools index {output.bam}
~~~

~~~
# ONT reads
winnowmap -t 48 -W {genome}_repetitive_k15.txt -ax map-ont -y {assembly_fasta} {fastq_with_meth_calls} >
{output.sam}
samtools view -@ 24 -Sb -F 2048 {output.sam} | samtools sort -@ 24 -T {temporary_directory} - > {output.bam}
samtools index {output.bam}
~~~

Aggregated methylation percentages at all CpGs were obtained using modbam2bed (v0.10.0, https://github.com/epi2me-labs/modbam2bed) with bases with >0.8 probability called “methylated” and bases with <0.2 probability called “unmethylated”:

~~~
modbam2bed -t 48 -e -m 5mC --cpg -a 0.20 -b 0.80 {assembly_fasta} {output.bam} > {output.bed}
~~~

### CUT&Tag library preparation

For anti-CENP-A CUT&Tag library preparation was used the CUT&Tag-IT kit from Active Motif (Cat. No. 53160). Each experiment was performed for 500,000 fresh cells. Fresh cells were washed using 1X Wash Buffer and nuclei were isolated and incubated with activated concanavalin A-coated magnetic beads in 2 mL PCR tubes at room temperature for 10 min. A 1:100 dilution of primary antibody anti-CENP-A (Human) mAb (Cat. No. D115-3) in antibody buffer was added and nuclei were incubated overnight at 4°C. The next day tubes were incubated on a magnetic tube holder and supernatants were discarded. Secondary antibody (Rabbit anti-mouse) was diluted at 1:100 in Dig-Wash buffer and nuclei were incubated for 1 h on an orbital rotator at room temperature. Nuclei were washed three times in Dig-Wash and then incubated with a 1:100 dilution of CUT&Tag-IT pA–Tn5 Transposomes for 1 h on an orbital rotator at room temperature. After, 125 μl of tagmentation buffer was added to each sample. To stop tagmentation, 4.2 μl 0.5 M EDTA, 1.25 μl 10% SDS and 1.1 μl 10 mg ml^−1^ proteinase K was added to each reaction and incubated at 55 °C for 1 h. DNA was barcoded and amplified using the following conditions: a PCR mix of 25 μl NEBNext 2× mix, 2 μl each of barcoded forward and reverse 10 μM primers, and 21 μl of extracted DNA was amplified at: 58°C for 5 min, 72°C for 5 min, 98°C for 45 s, 16× 98°C for 15 s followed by 63°C for 10 s, 72°C for 1 min. Amplified DNA libraries were purified by adding a 1.1× volume of SPRI beads to each sample and incubating for 10 min at 23°C. Samples were placed on a magnet and unbound liquid was removed. Beads were rinsed twice with 80% ethanol, and DNA was eluted with 20 μl Elution buffer. All individually i7-barcoded libraries were mixed at equimolar proportions for sequencing.

### CUT&Tag libraries and sequencing

Libraries were quantified and individually converted to process on the Singular Genomics G4 with the SG Library Compatibility Kit (Cat. No. 700141), following the “Adapting Libraries for the G4 – Retaining Original Indices” protocol. The converted libraries were sequenced in individual lanes on an F3 flow cell (Cat. No. 700125) on the G4 instrument, using Instrument Control Software 23.08.1-1 with 100 bp paired-reads. Following sequencing, sgdemux 1.2.0 was run to generate FASTQ files.

### CUT&Tag bioinformatic analysis

CUT&Tag sequencing reads were trimmed using the trim-galore tool (v0.6.10, https://github.com/FelixKrueger/TrimGalore), which included adapter removal. The trimmed reads of each sample were then aligned to the corresponding generated *de novo* assemblies using bowtie2 (v2.5.3)^14^. Post-alignment, the reads were sorted and indexed using samtools (v1.17)^13^, to then extract depth information for primary alignments with mosdepth^15^.

### Pairwise sequence identity heatmaps

To generate pairwise sequence identity heatmaps of each centromeric region, we used a modified version of StainedGlass (v0.6)^16^ with the following parameters: window=5000, mm_f=30000, and mm_s=1000. Our modifications were applied to visualize the identity heatmaps with methylation and CENP-A CUT&Tag information included at the bottom.

### Synteny plots

To visualize the alignment between the generated assemblies and CHM13, we used NGenomeSyn^17^ to generate the synteny plots, which were then manually curated.

### Hi-C data analysis

Hi-C reads were mapped to CHM13 with the BWA aligner^18^, configured to handle the chimeric nature of Hi-C reads by allowing local mapping and tuning the parameters to minimize gaps. Following read mapping, we constructed the Hi-C contact matrix by specifying a bin size of 10,000 bp and incorporating restriction site information using HiCExplorer tools^19^. The resulting matrices were then binned at various resolutions (100kb, 200kb, 500kb) and corrected to normalize the contact frequencies across bins and remove GC and open chromatin biases. Finally, we visualized the corrected matrices using hicPlotMatrix, applying log transformation to handle the wide range of contact counts.

### Genome versions used

We leveraged multiple reference genomes and assemblies. The primary reference was T2T-CHM13v2.0. We also incorporated the recent diploid T2T-HG002v1.1 genome and 72 samples from the Human Pangenome Reference Consortium (HPRC). The HPRC samples were assembled using Verkko v2.1^1^, employing a combination of sequencing technologies for each sample. The assembly process utilized PacBio High-Fidelity (HiFi) reads and Oxford Nanopore Technology (ONT) long reads. For phasing, we primarily used short Illumina reads. In cases where trio information was unavailable, Hi-C reads were used for phasing instead.

