## Supplementary figures 1-14 for "The formation and propagation of human Robertsonian chromosomes"

#### Supplementary Figure 1. Assembly graphs of the t(14;21)-bearing GM03417 cell line

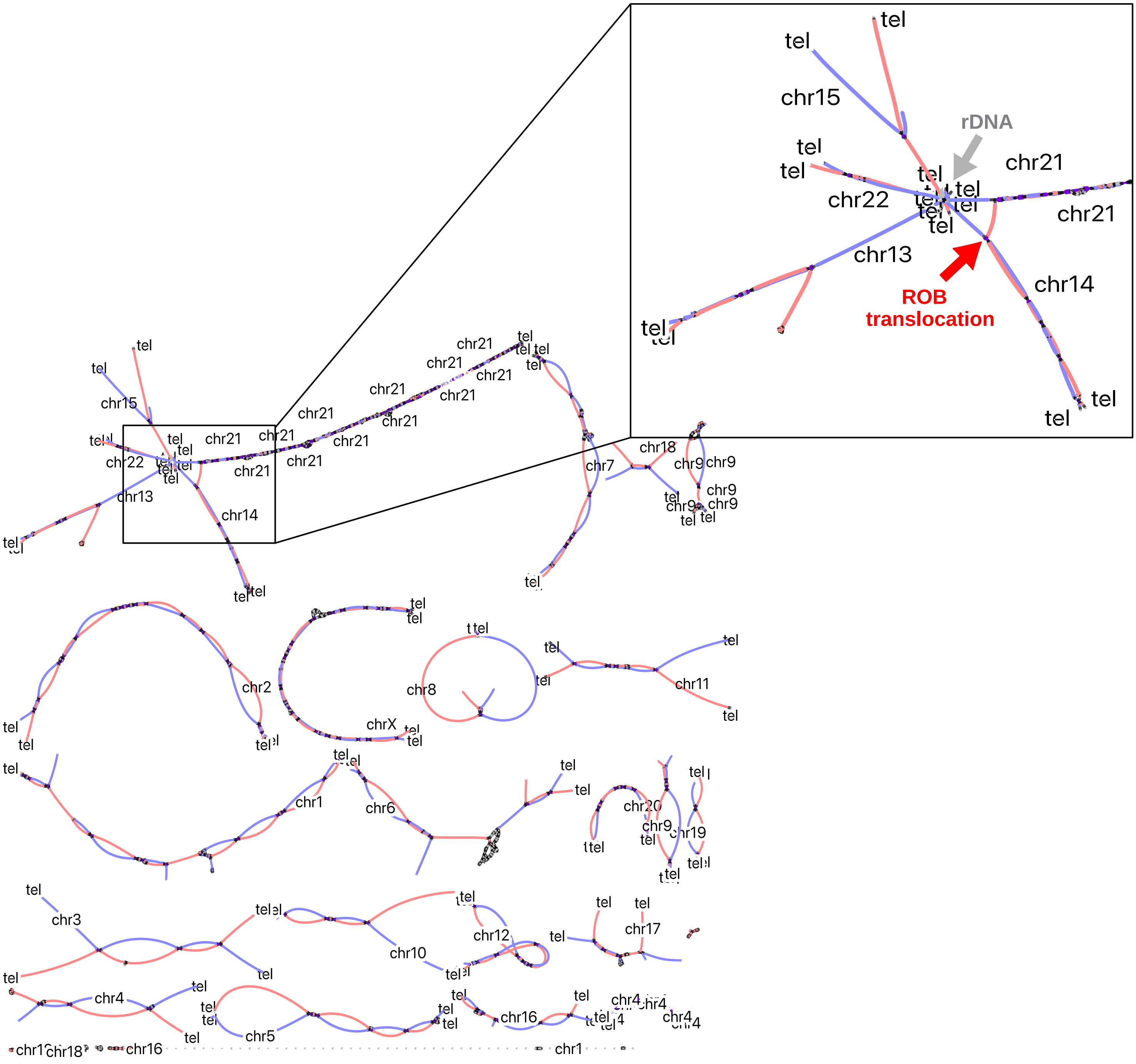


Assembly graph of GM03417 integrating PacBio and ONT reads visualized using Bandage (https://doi.org/10.1093/bioinformatics/btv383). Each chromosome is mostly resolved as a single connected component, with the exception of the acrocentric chromosomes (13, 14, 15, 21 and 22), which are joined by the highly similar rDNA arrays. The graph is colored by Hi-C information used to phase both haplotypes (haplotype 1, red; haplotype 2, blue). The inset shows a portion of the graph involving the acrocentric chromosomes, highlighting the graph node representing the Robertsonian translocation, which connects chromosomes 14 and 21.

#### Supplementary Figure 2. Assembly graphs of the t(13;14)-bearing GM03786 cell line

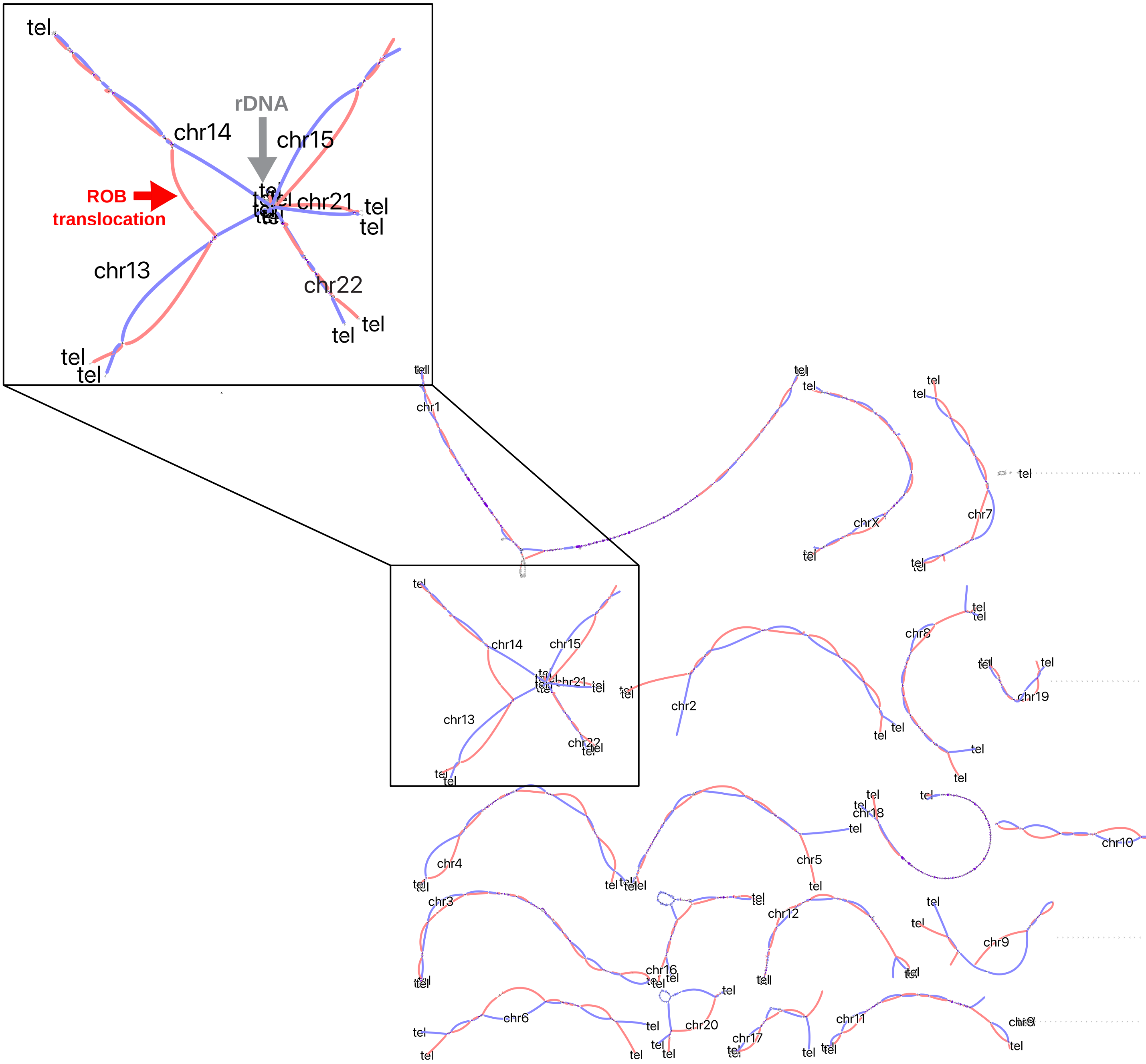


Assembly graph of GM03786 integrating PacBio and ONT reads visualized using Bandage. Each chromosome is mostly resolved as a single connected component, with the exception of the acrocentric chromosomes (13, 14, 15, 21 and 22), which are joined by the highly similar rDNA arrays. The graph is colored by Hi-C information used to phase both haplotypes (haplotype 1, red; haplotype 2, blue). The inset shows a portion of the graph involving the acrocentric chromosomes, highlighting the graph node representing the Robertsonian translocation, which connects chromosomes 13 and 14.

#### Supplementary Figure 3. Assembly graphs of the t(13;14)-bearing GM04890 cell line

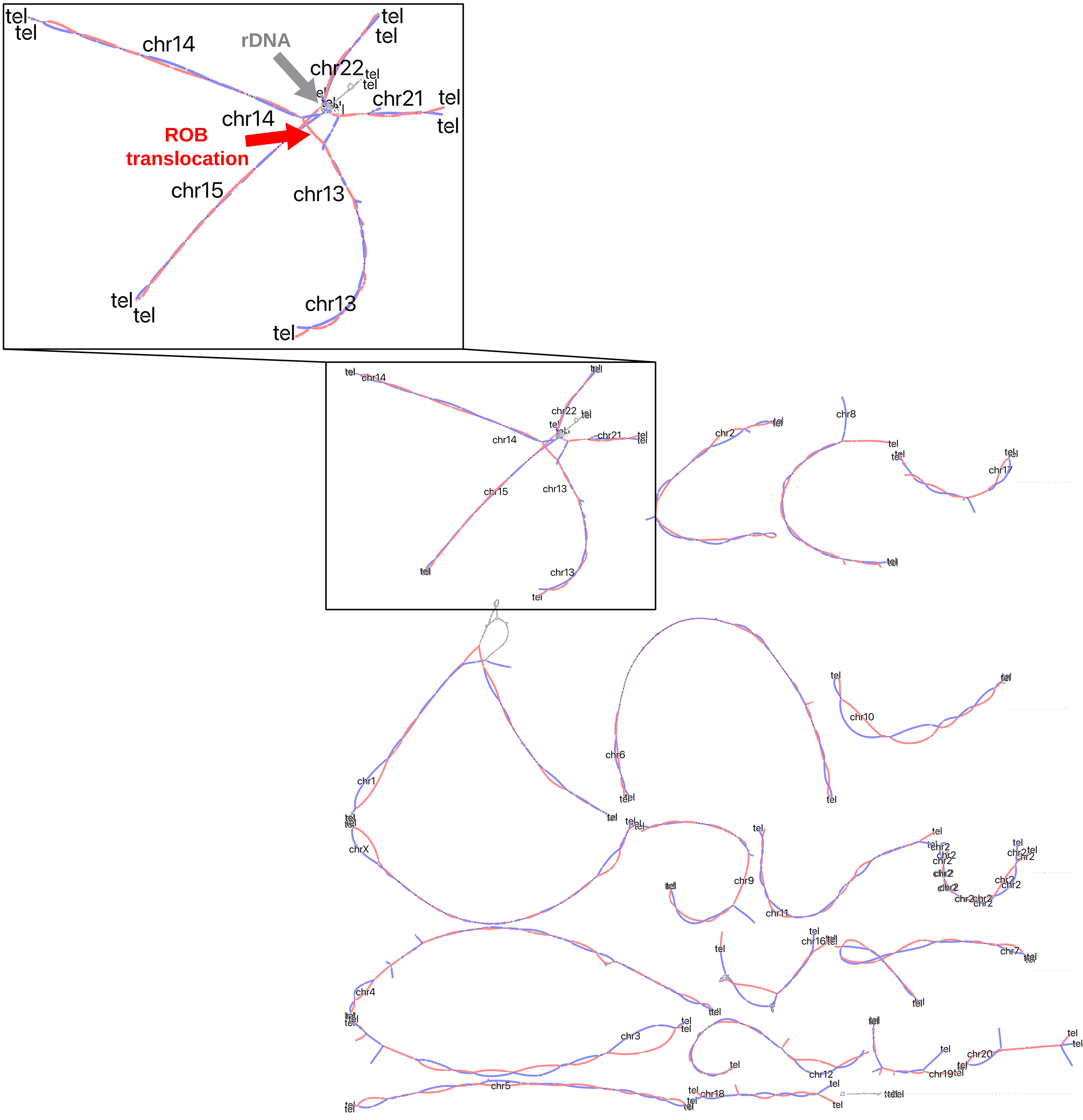


Assembly graph of GM04890 integrating PacBio and ONT reads visualized using Bandage. Each chromosome is mostly resolved as a single connected component, with the exception of the acrocentric chromosomes (13, 14, 15, 21 and 22), which are joined by the highly similar rDNA arrays. The graph is colored by Hi-C information used to phase both haplotypes (haplotype 1, red; haplotype 2, blue). The inset shows a portion of the graph involving the acrocentric chromosomes, highlighting the graph node representing the Robertsonian translocation, which connects chromosomes 13 and 14.

#### Supplementary Figure 4. Assembly gene completeness evaluation

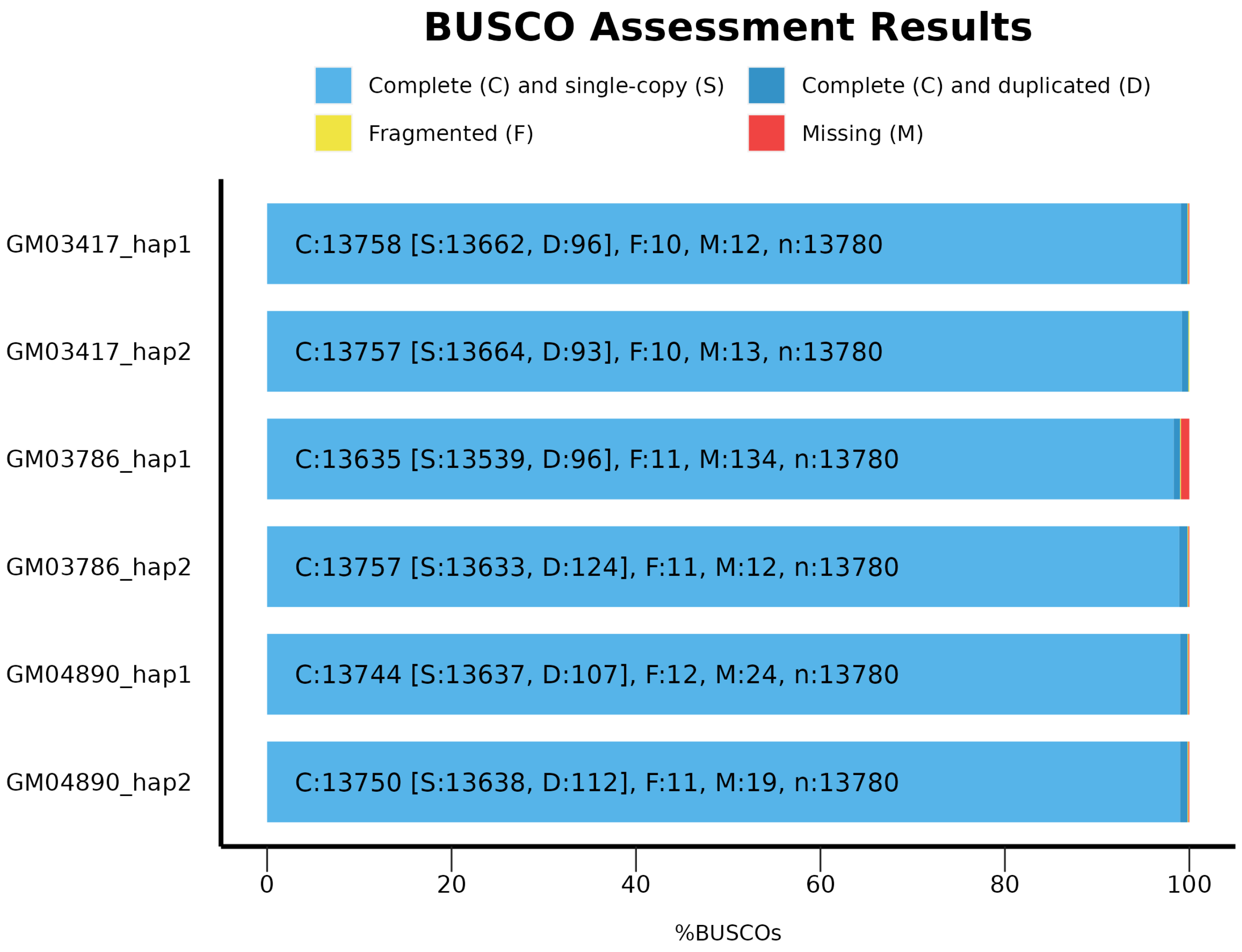


BUSCO (Benchmarking Universal Single-Copy Orthologs) assessment results for the diploid genome assemblies of the three ROB-bearing cell lines analyzed (GM03417, GM03786, and GM04890), with two haplotypes each (hap1 and hap2). The horizontal bars show the percentage of BUSCOs found in each assembly, categorized as follows: Light blue: Complete (C) and single-copy (S) BUSCOs; Dark blue: Complete (C) and duplicated (D) BUSCOs; Yellow: Fragmented (F) BUSCOs; Red: Missing (M) BUSCOs. Each bar is annotated with the exact counts for each category, along with the total number of BUSCOs assessed (n:13780, primates set).

#### Supplementary Figure 5. Coverage analysis of Robertsonian translocation breakpoints

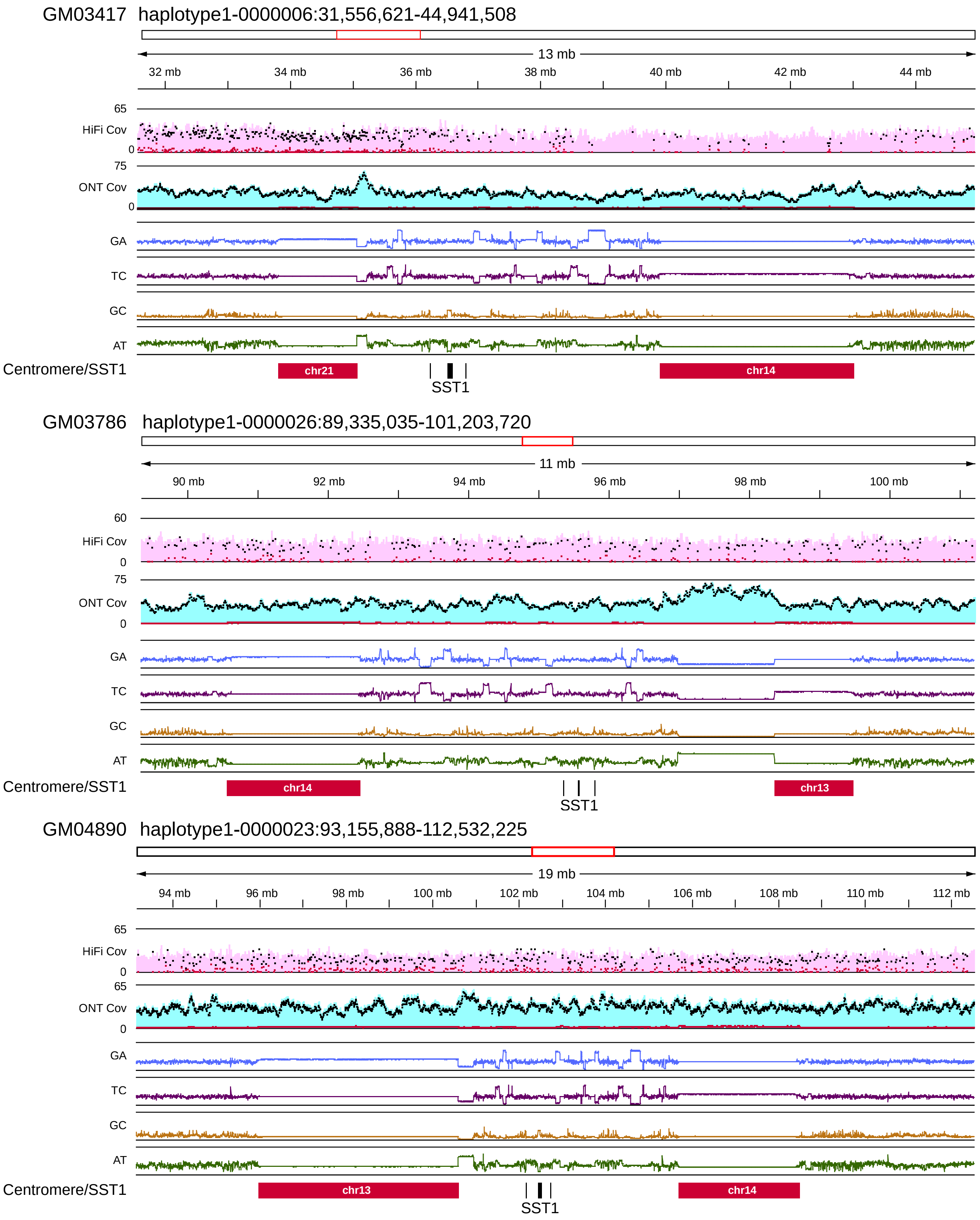


HiFi and ONT read alignments for the ROB-bearing cell lines analyzed (GM03417, GM03786, GM04890) are shown. Tracks from top to bottom: genomic coordinates, HiFi coverage (black: primary allele, red: secondary allele), ONT coverage, and nucleotide content (GA, TC, GC, AT). Annotations below indicate centromeric arrays (red bars) and SST1 arrays (black bars). The consistent coverage across breakpoints supports the accuracy of the assemblies.

#### Supplementary Figure 6. Impact of ROBs on chromosome arm interactions

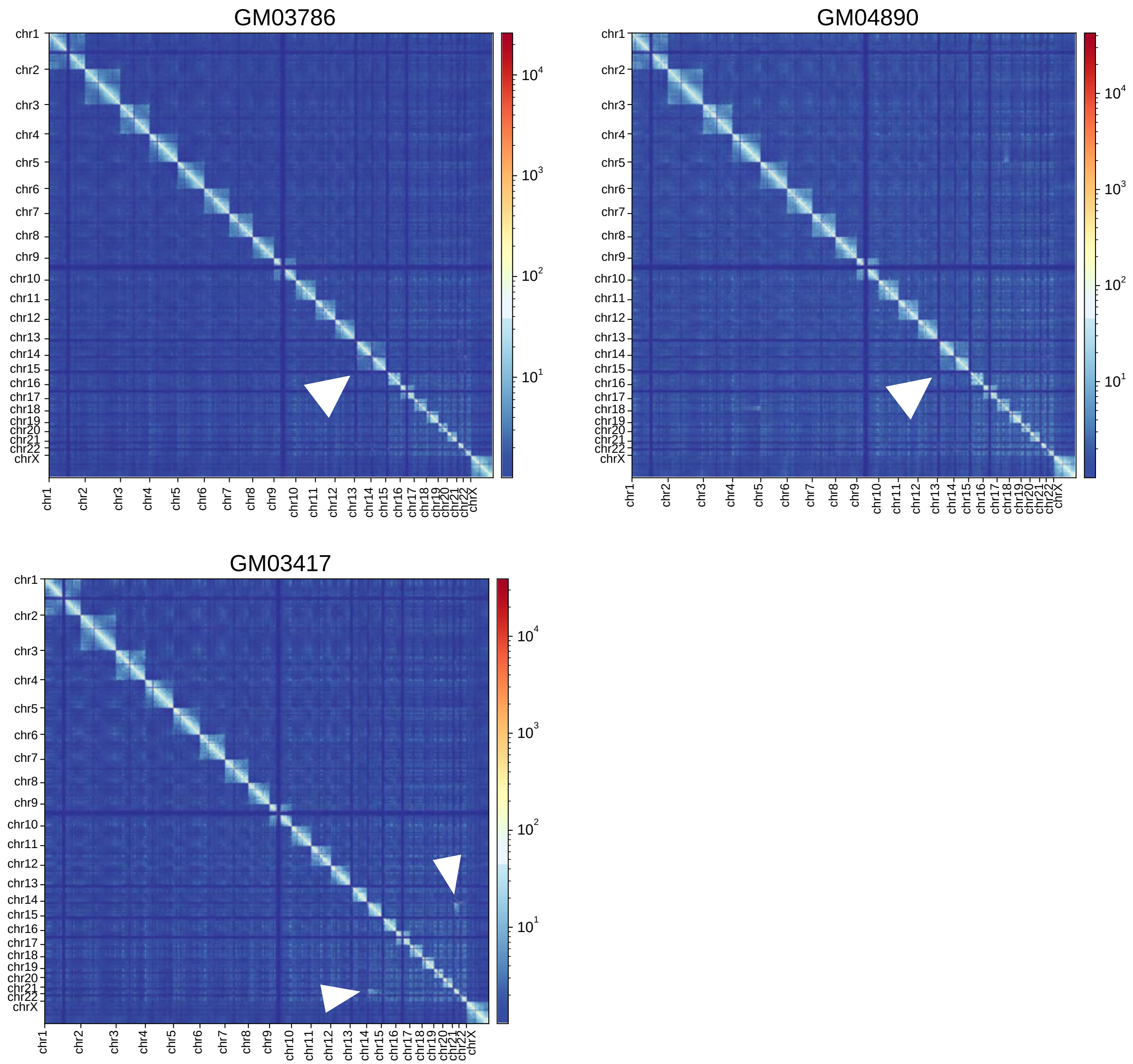


Genome-wide Hi-C interaction matrices (with a resolution of 200 kb) for the three ROB-bearing cell lines analyzed. Chromosomes involved in the Robersonian event show high interchromosomal interactions (arrowheads).

#### Supplementary Figure 7. Cytogenetic evidence for the structural changes in ROBs

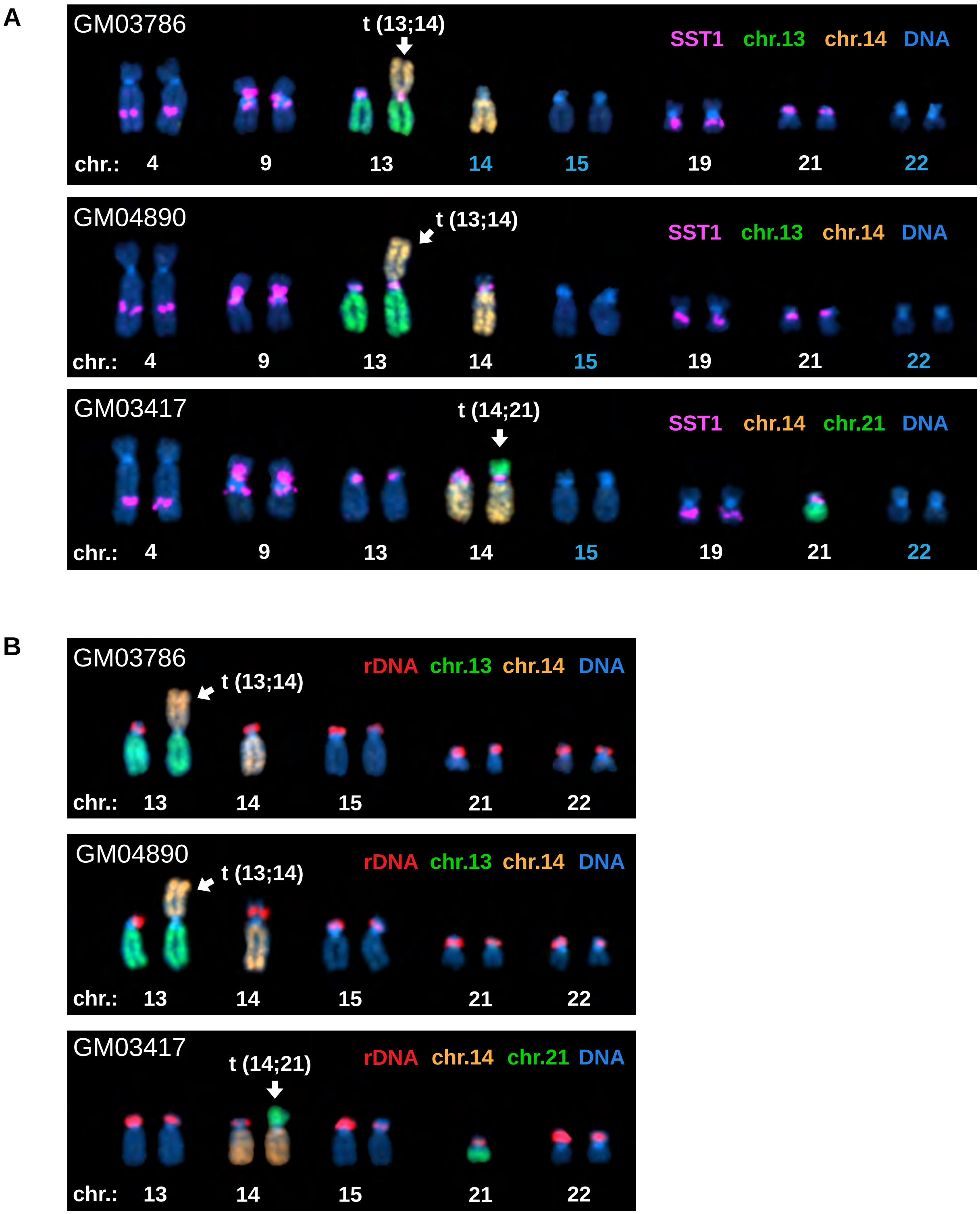


Fluorescence in situ hybridization (FISH) was used to visualize the arrangement of SST1 and 45S rDNA arrays on acrocentric chromosomes in the ROB-bearing cell lines. **A.** Extended karyograms of GM03786, GM04890, and GM03417 cell lines labeled by FISH with SST1 probe (magenta) and whole chromosome paints (14 in orange, 13 and 21 in green). All acrocentric chromosomes and other chromosomes with detectable SST1 signals are shown. Acrocentric chromosomes lacking SST1 signal are denoted with blue numbers. DNA was counter-stained with DAPI. **B.** Extended karyograms of acrocentric chromosomes only labeled by FISH with rDNA probe (red), whole chromosome paints, and DAPI. Note lack of rDNA signal on ROBs in all cell lines.

#### Supplementary Figure 8. Schematic representation of common ROB formation

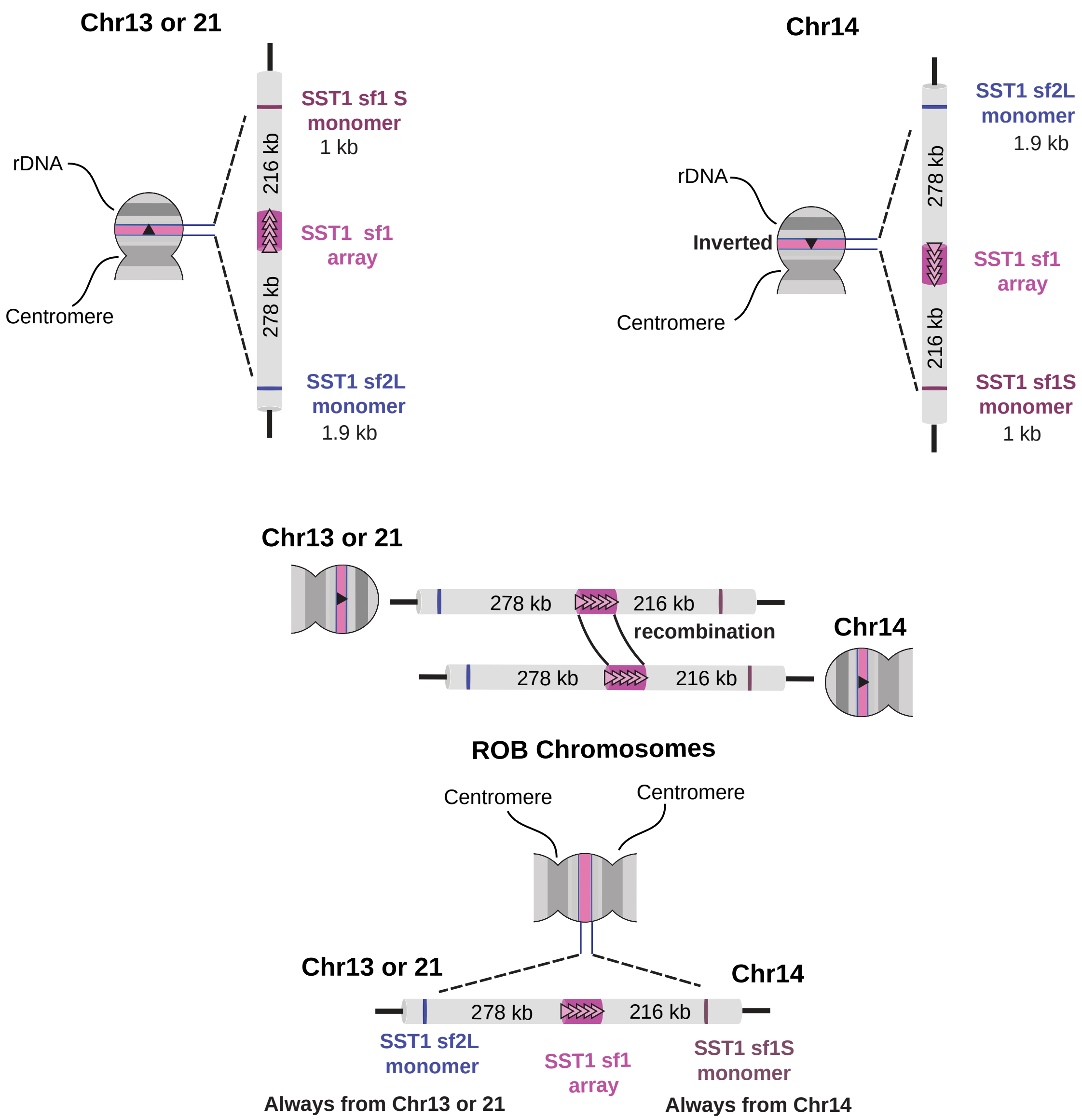


Top panel: Structure of the short arms of chromosomes 13/21 and chromosome 14 before fusion. Centromeric, rDNA, and SST1 arrays are highlighted. Note the inverted orientation of the SST1 type 1 array on 14 relative to 13/21. Middle panel: Alignment of 13/21 and 14 during meiosis, showing potential recombination within the SST1 type 1 array region. Bottom panel: Resulting ROB structure after fusion. The ROB retains two centromeres and the SST1 arrays from both parent chromosomes. The central multicopy SST1 array is flanked by sequences from 13 and 14 or 14 and 21. The fusion is within the SST1 subfamily 1 array. The size of the array is variable. The SST1 type 2L monomer is derived from 13 or 21, while the SST1 type 2S monomer is from 14. The rDNA regions are lost in the fusion process. Key features are color-coded: magenta for SST1 type 1 array, pink for SST1 type 2S monomer, blue for SST1 type 2L monomer, and gray for centromeres and rDNA regions. Distances between features are indicated in kilobases (kb).

**Supplementary Figure 9. Sequence alignment comparison among SST1 monomer consensuses from each chromosome in CHM13 with a major array**

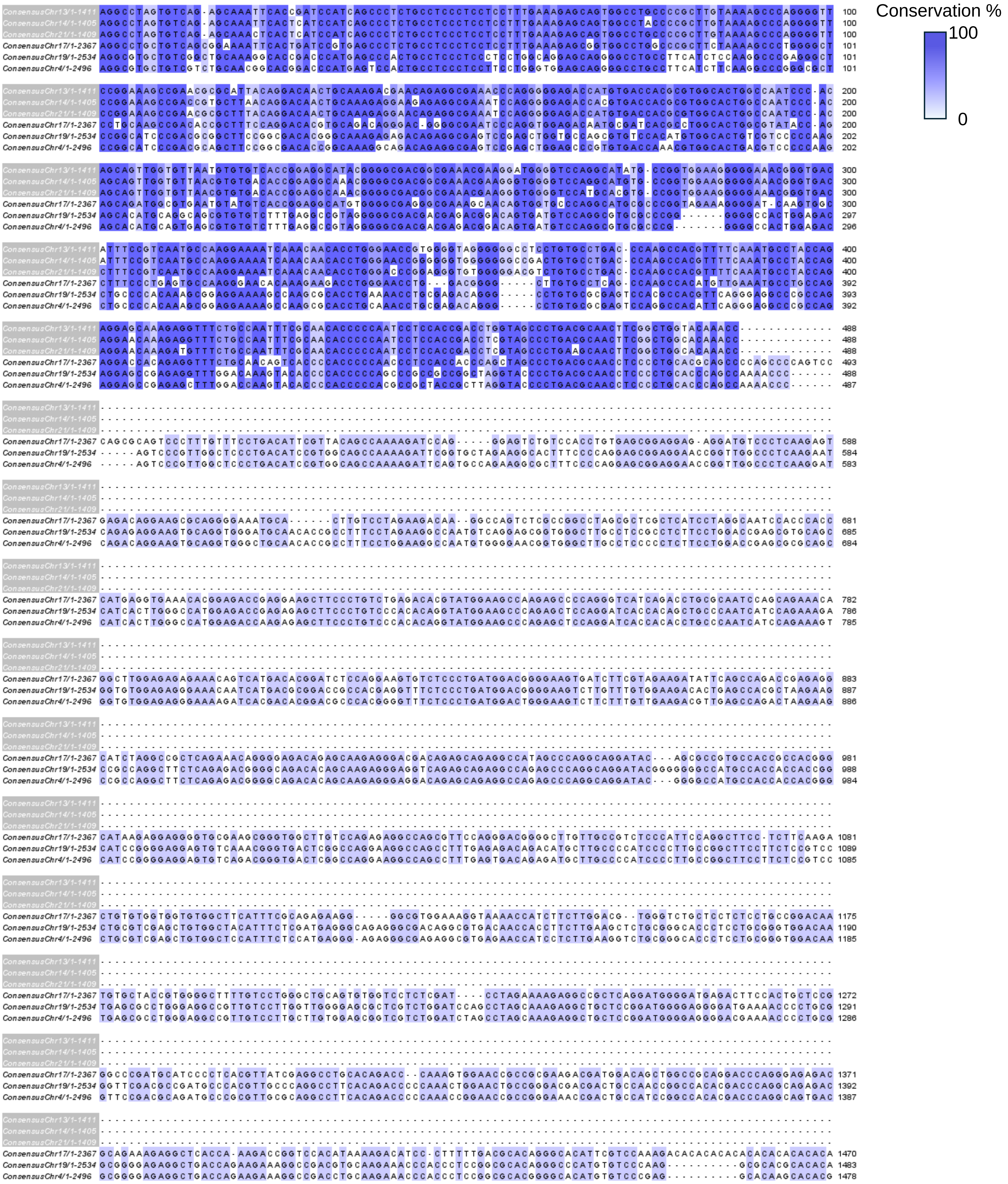

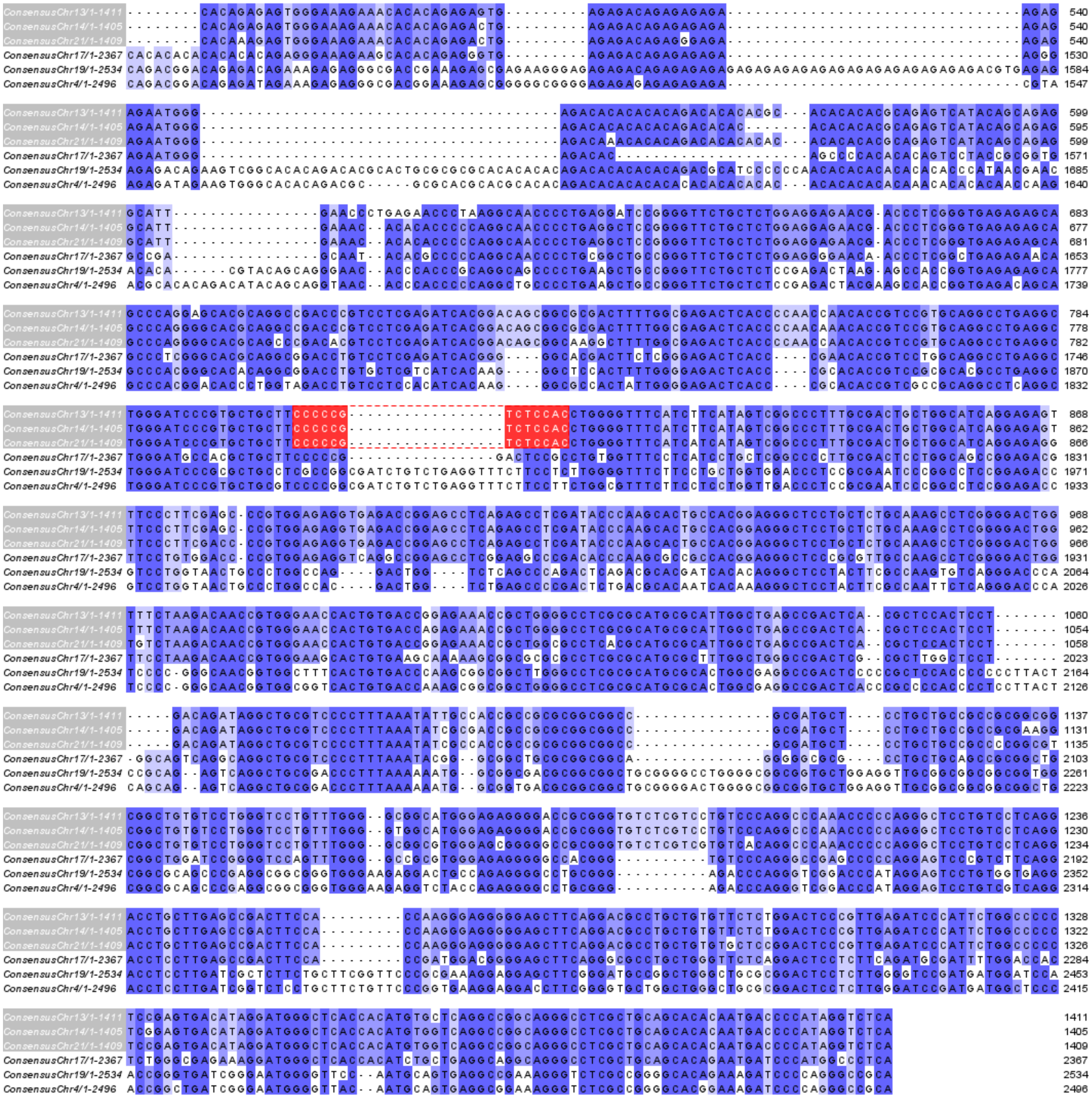

Each row corresponds to the consensus sequence for monomers of SST1 derived from major arrays on chromosomes 13, 14, 21, 17, 19, and 4. The consensuses from chromosomes 13, 14, and 21 (gray labels) have in common a large gap in the middle of the macrosatellite as well as a predicted PRDM9 DNA binding site (red). Darker shades indicate more conservation.

#### Supplementary Figure 10. Representative karyogram showing chromosomal location of SST1 SF1 arrays mapped by FISH.

A

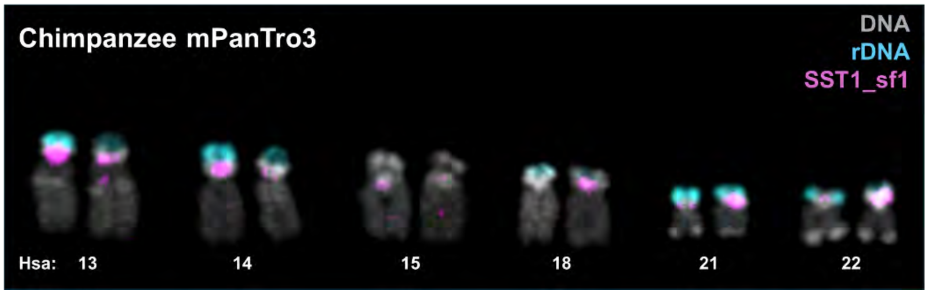


B

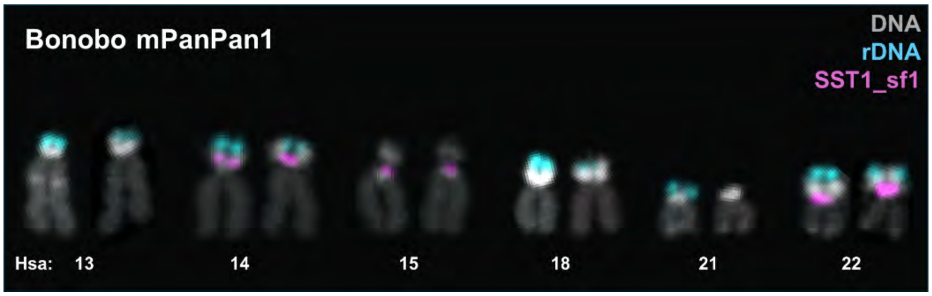


Differential and shared locations of SST1 sf1 arrays in the p arm of acrocentric chromosomes in **(A)** chimpanzee and **(B)** bonobo. FISH was performed on chromosome spreads from cultured fibroblasts. The panels show SST1 sf1 (magenta) and 45S rDNA (cyan) probes with DAPI staining (gray).

### Supplementary Figure 11. Phylogenetic clustering of segmental duplications involving SST1 in nonhuman primates.

A

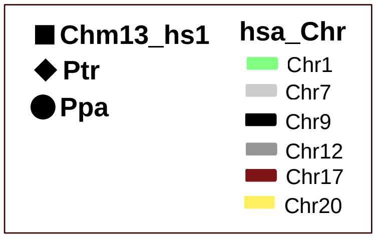


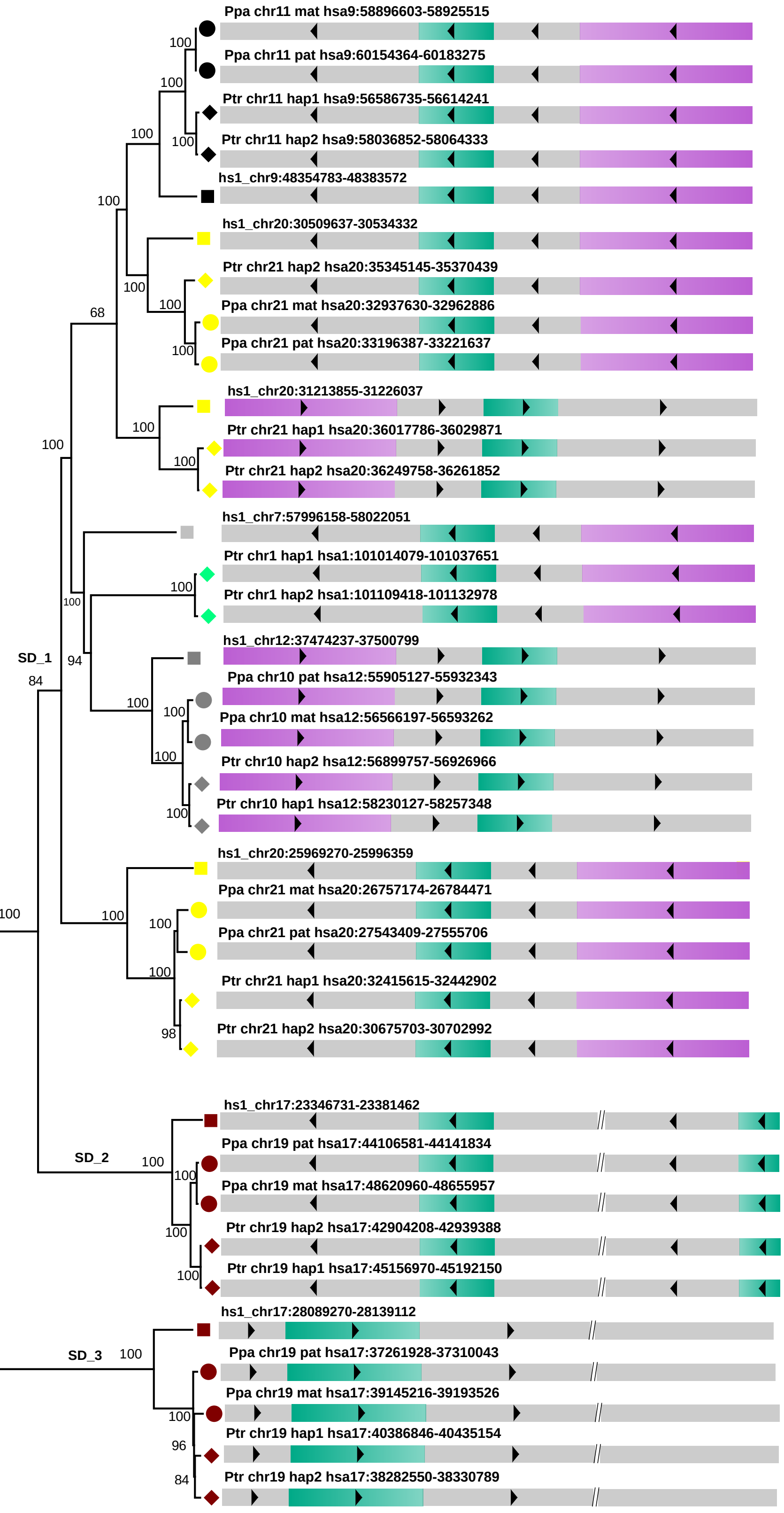


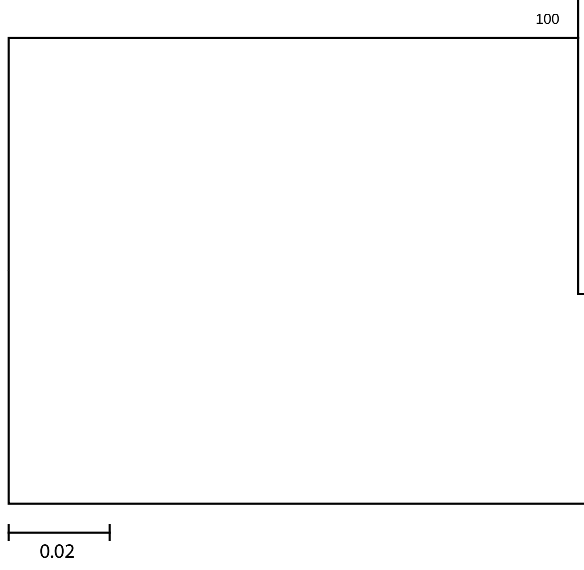


**A)** Three types of segmental duplications (SD1, SD2, SD3) ranging in size from ~28-50 kb that included SST1 were identified in human (CHM13, squares), chimpanzee (diamonds), and bonobo (circles) genomes. Maximum likelihood phylogenetic reconstruction was performed using the GTR +Gamma substitution model with bootstrap values for 1000 iterations. Each chromosome associated with the segmental duplication event is indicated in the color key. B) All three segmental duplication structures contain Y-like SST1 sequences (sf3, green), as well as Y-like alpha-satellite sequences (grey), suggesting the region originated from a Y chromosome in the common ancestor.

B

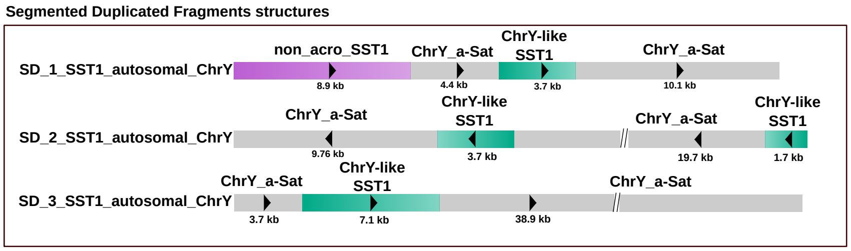


**B)** All three segmental duplication structures contain Y-like SST1 sequences (sf3, green), as well as Y-like alpha-satellite sequences (grey), suggesting the region originated from a Y chromosome in the common ancestor.

**Supplementary Figure 12. CENP-B localization on dicentric ROBs**

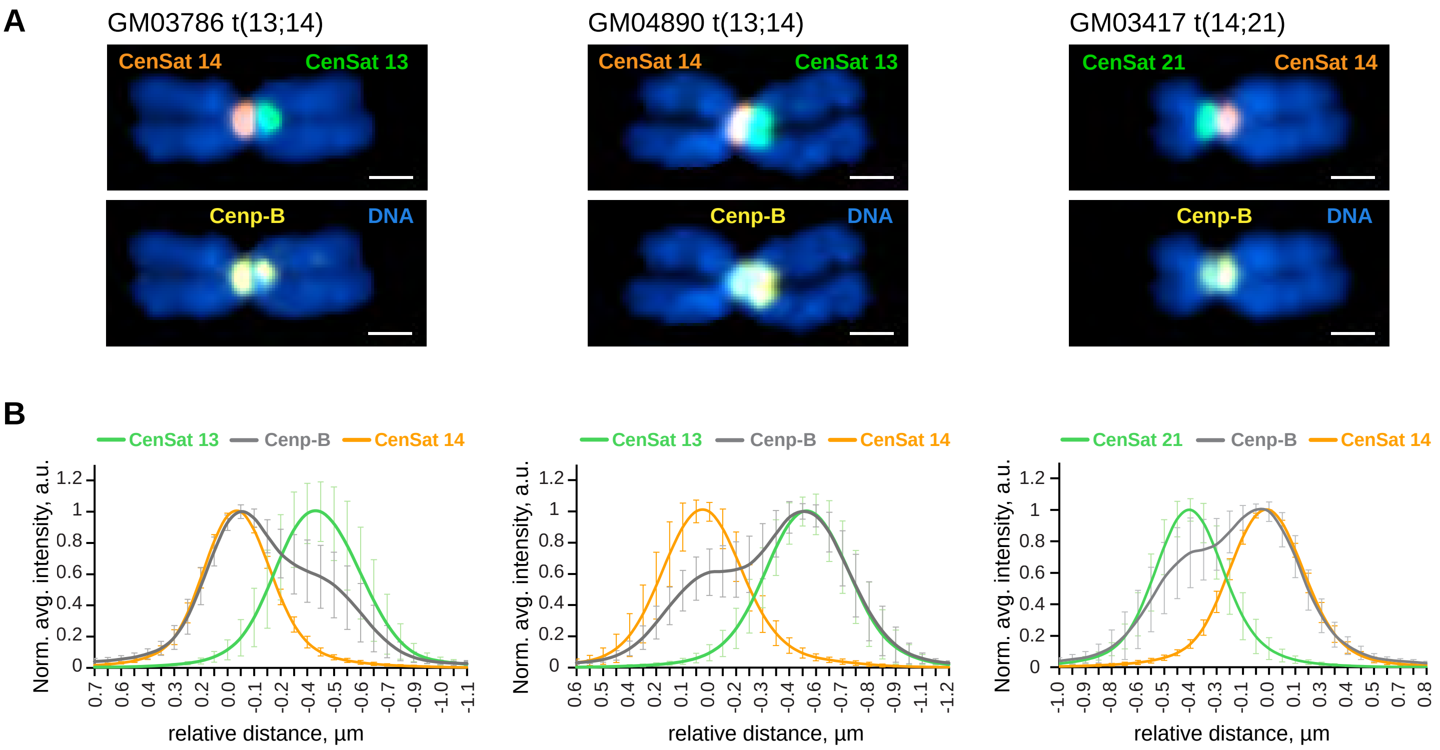


**A)**. Representative images of ROBs from GM03786, GM04890, and GM03417 cell lines labeled by immuno-FISH with centromeric satellite probes and antibody against CENP-B. Top panels display chromosomes labeled with centromeric satellite probes for CenSat 14/22 (orange) and CenSat 13/21 (green). Bottom panels show the same chromosomes with CENP-B labeling (yellow). DNA was counter-stained with DAPI (blue). Note the CENP-B localization on both centromeres. Scale bar is 1 µm. **B)** Averaged intensity profiles of lines drawn through the centromeres of at least 10 chromosomes. Intensity profiles were aligned to the peak of the Gaussian of the CenSat 14 signal and normalized to the maximum intensity of each channel. Error bars denote standard deviations.

#### Supplementary Figure 13. Schematic of centromeric arrays on the ROBs

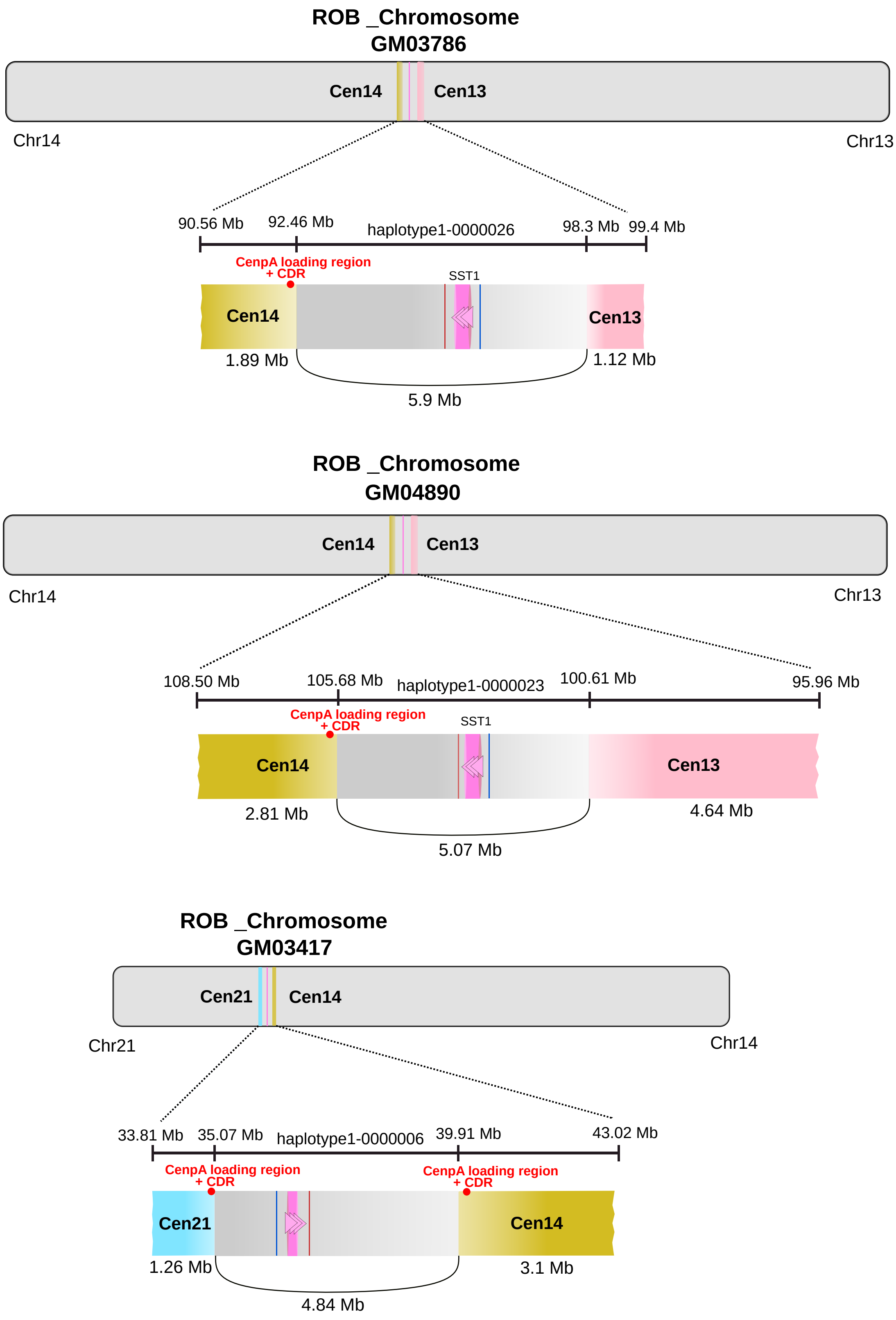


The size (in Mb) and arrangement of the two centromeric arrays are shown for each of the three ROBs. The distance between centromeric arrays ranges from 4.8-5.9 Mb. The position of CENP-A, as determined by CUT&Tag, and the dip in methylation at the centromere (CDR), based on ONT reads, are both indicated (red).

### Supplementary Figure 14. Imaging and genomic analysis of chromosomes 13, 14, and 21 in assembled genomes

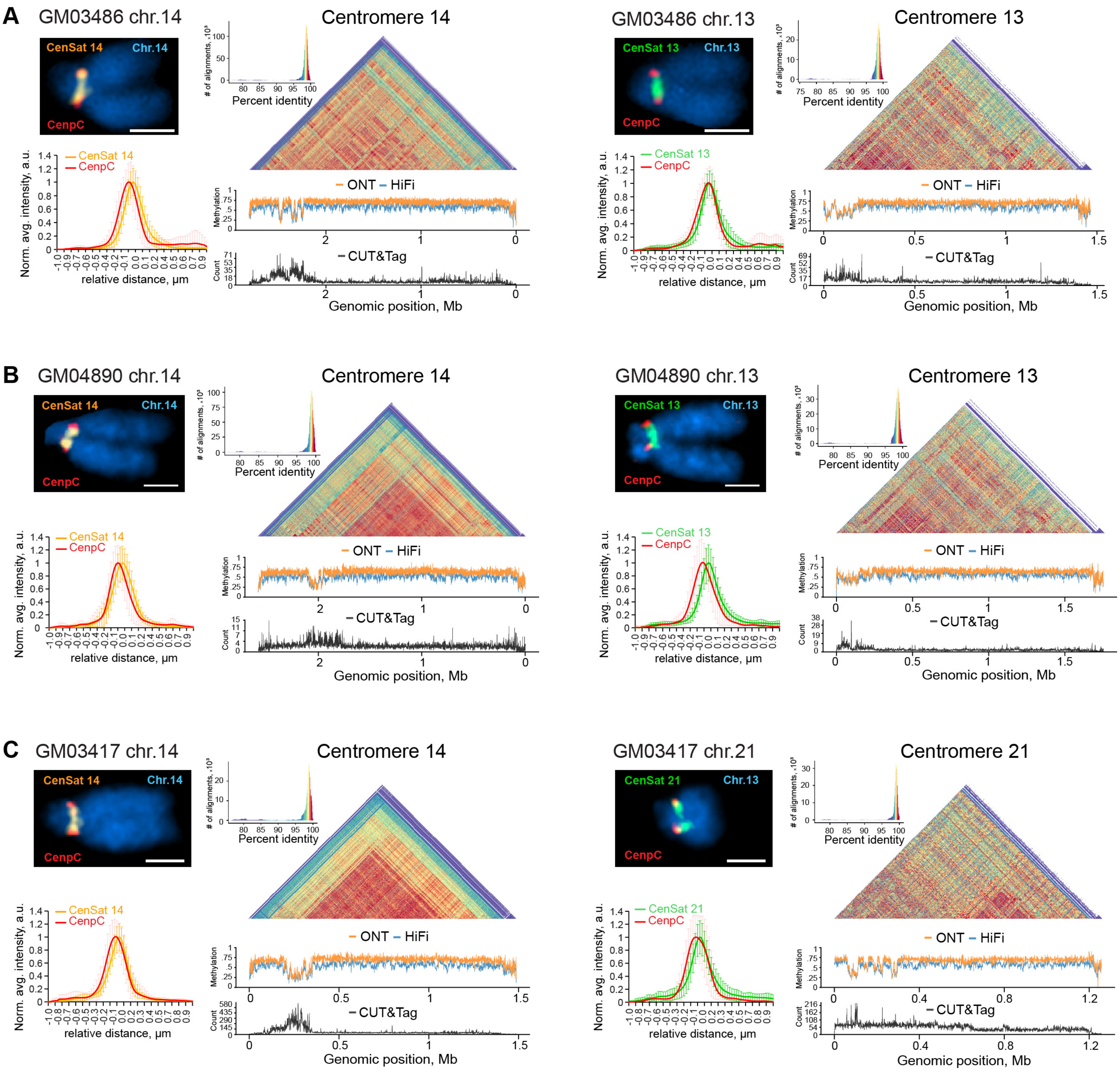


ImmunoFISH, DNA methylation, and CENP-A CUT&Tag analysis of normal copies of acrocentric chromosomes from **(A)** GM03786, **(B)** GM04890 and **(C)** GM03417 cell lines are shown. The left panels show representative structured illumination super-resolution images of normal acrocentric chromosomes labeled by immuno-FISH with centromeric satellite probes for CenSat 14/22 (orange), CenSat 13/21 (green), and anti-CENP-C antibody (red). DNA was counterstained with DAPI. Scale bar is 1 µm. Note single CENP-C foci on corresponding centromeres. Plots below show averaged intensity profiles of at least 10 lines drawn through the centromeric regions of sister chromatids. Intensity profiles were aligned to the peak of the Gaussian of the corresponding CenSat signals and normalized to the maximum intensity of each channel. Error bars denote standard deviations. The right panels display corresponding heatmaps of sequence similarity calculated for 5 kb bins for each centromere. Below the heatmaps, DNA methylation tracks show methylation calls from ONT (orange) or PacBio HiFi (blue) sequencing. Hypomethylated regions correspond to CENP-A peaks on CUT&Tag tracks below (black), indicating active centromere localization.
